# Proteostasis and lysosomal repair deficits in transdifferentiated neurons of Alzheimer’s disease

**DOI:** 10.1101/2023.03.27.534444

**Authors:** Ching-Chieh Chou, Ryan Vest, Miguel A. Prado, Joshua Wilson-Grady, Joao A. Paulo, Yohei Shibuya, Patricia Moran-Losada, Ting-Ting Lee, Jian Luo, Steven P. Gygi, Jeffery W. Kelly, Daniel Finley, Marius Wernig, Tony Wyss-Coray, Judith Frydman

**Author notes:** These authors jointly supervised this work: Ching-Chieh Chou, Judith Frydman.

## Abstract

Aging is the most prominent risk factor for Alzheimer’s disease (AD). However, the cellular mechanisms linking neuronal proteostasis decline to the characteristic aberrant protein deposits in AD brains remain elusive. Here, we develop transdifferentiated neurons (tNeurons) from human dermal fibroblasts as a neuronal model that retains aging hallmarks and exhibits AD-linked vulnerabilities. Remarkably, AD tNeurons accumulate proteotoxic deposits, including phospho-Tau and Aβ, resembling those in AD patient and APP mouse brains. Quantitative tNeuron proteomics identify aging and AD-linked deficits in proteostasis and organelle homeostasis, most notably in endosome-lysosomal components. Lysosomal deficits in aged tNeurons, including constitutive lysosomal damage and ESCRT-mediated lysosomal repair defects, are exacerbated in AD tNeurons and linked to inflammatory cytokine secretion and cell death. Supporting lysosomal deficits’ centrality in AD, compounds ameliorating lysosomal function reduce Aβ deposits and cytokine secretion. Thus, the tNeuron model system reveals impaired lysosomal homeostasis as an early event of aging and AD.

## Introduction

Aging is central to Alzheimer’s disease (AD) and linked to a decline in protein homeostasis (proteostasis) and organelle homeostasis^1, 2^, including the endosome-lysosome^3–6^. The mechanistic underpinnings of these defects during human brain aging and disease remain poorly understood. Genetic models for AD that accumulate Aβ or tau aggregates show endosome-lysosomal dysfunction^2, 7–9^. It is a long-standing hypothesis that AD pathologies are mediated by non-cell autonomous effects whereby extra-cellular aggregates taken up by neurons lead to lysosomal damage and cell death^10–13^. However, this hypothesis does not explain early disease events leading to initial dysfunction, highlighting the unmet need for patient neuronal models to better understand the molecular origins of aging and AD pathological processes.

Developing cellular systems to study proteostasis and organellar phenotypes caused by aging and AD in human neurons remains challenging. While human neurons from postmortem brains are widely studied at single-cell transcriptomics^14^, changes in the cytonuclear and organellar proteostasis networks are often not apparent from these datasets. Another common tool are induced pluripotent stem (iPS) cells and the derived lineages^15^. However, this process restores youthfulness to the induced neurons (iNeurons), forgoing the key contribution of aging to neurodegeneration^15, 16^. The recent development of transdifferentiated neurons (herein tNeurons) directly from human adult somatic cells enables studying neurodegenerative diseases while maintaining the contribution of aging^16, 17^. tNeurons retain aging and disease phenotypes ^18–21^ but due to limiting samples, they are primarily examined via transcriptomics, which do not generally reflect proteome- and organelle-wide changes. We improved the transdifferentiation approach to generate tNeurons from dermal fibroblasts and performed quantitative proteomic analyses combined with biochemical and functional analyses comparing tNeurons obtained from healthy young and aged donors, as well as from patients with sporadic AD (sAD) and familial AD (fAD). We find that aging and AD create a cell-autonomous vulnerable state in neurons characterized by constitutive lysosomal damage, impaired proteostasis and defective repair of compromised lysosomes leading to intra-neuronal protein deposition and secretion of inflammatory cytokines. Our findings may lead to potential therapeutic strategies against aging and AD.

## Results

### Aging and proteostasis signatures in fibroblasts and tNeurons

We harnessed transcription factors *Brn2*, *Ascl1*, *Myt1l*, and *Ngn2* along with small molecules to transdifferentiate human fibroblasts into cortical neurons (Fig. 1a). Fibroblasts were collected from 8 healthy young (age: 25.6±4.9) and 12 aged donors (age: 70.3±5.9), as well as 16 aged patients with sAD (herein aged/sAD, age: 70.4±9.2). Fibroblasts from 5 middle-aged fAD patients carrying *PSEN1* mutations (herein fAD-PSEN1, age: 47.2±10.2) were subsequently derived for certain experiments (Supplementary Table 1). Fibroblasts from aged and aged/sAD donors showed an increase in DNA damage measured by γ-H2AX puncta (Fig. 1b) and a global loss of epigenetic marker, H3K9me3 (Fig. 1c). We observed H4K16ac enriched during aging but reduced with AD (Fig. 1c), recapitulating findings in human brain^22^.

**Fig. 1.**
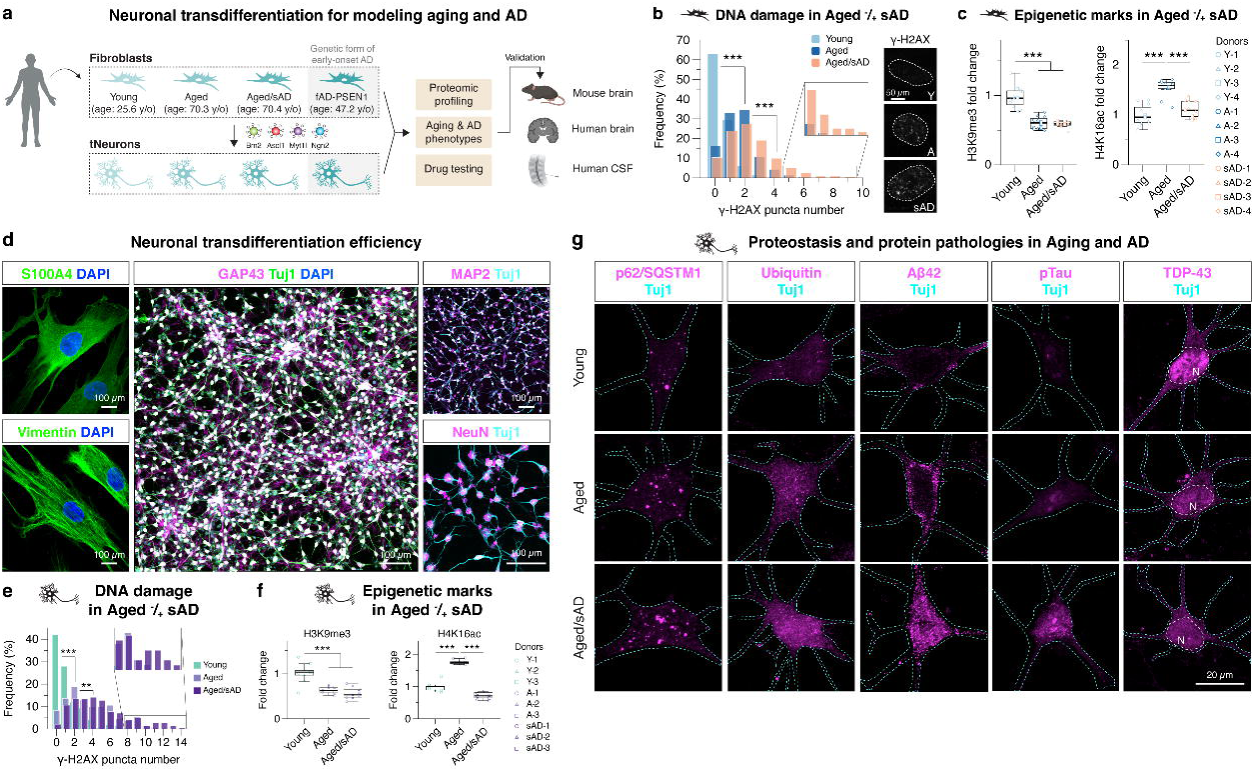
Transdifferentiating human adult fibroblasts into neurons reveals signatures of aging and Alzheimer’s disease (AD). **(a)** Human dermal fibroblasts are collected from donors of healthy young and aged, aged with sporadic AD (aged/sAD) and familial AD with *PSEN1* mutations (fAD-PSEN1). Fibroblasts and the transdifferentiated neurons (tNeurons) are used for a variety of experiments and the findings are validated in post-mortem brain tissue and cerebrospinal fluid (CSF). **(b)** Levels of DNA damage measured by numbers of the nuclear foci of γ-H2AX immunofluorescence (IF) in human fibroblasts. *n* = 248 (young), 304 (aged) and 252 (aged/sAD) cells from three donors and three independent experiments. Scale bar: 50 μm. **(c)** Age- and AD-related epigenetic alterations measured by histone modifications H3K9me3 and H4K16ac IF in human fibroblasts. H3K9me3: *n* = 252 (young), 241 (aged) and 237 (aged/sAD) cells from four donors and three independent experiments; H4K16ac: *n* = 157 (young), 167 (aged) and 141 (aged/sAD) cells from four donors and three independent experiments. **(d)** Representative images of human fibroblasts immunostained for S100A4 and Vimentin, and tNeurons immunostained for Tuj1, GAP43, MAP2 and NeuN with DAPI counterstaining at post-induction day (PID) 35. Scale bar: 100 μm. **(e)** IF quantification of DNA damage in tNeurons revealed by γ-H2AX. *n* = 150 (young), 132 (aged) and 141 (aged/sAD) cells from three donors and three independent experiments. **(f)** IF quantification of H3K9me3 and H4K16ac changes in tNeurons. H3K9me3: *n* = 95 (young), 123 (aged) and 124 (aged/sAD) cells from three donors and three independent experiments; H4K16ac: *n* = 112 (young), 115 (aged) and 117 (aged/sAD) cells from three donors and three independent experiments. **(g)** Representative images of proteostasis- and disease-associated protein markers in tNeurons, including autophagy adaptor p62/SQSTM1, ubiquitin, Aβ42, hyper-phosphorylated tau (pTau) and TDP-43. Cyan dash line outlines tNeuron morphology determined by Tuj1 staining. White dash line represents the nuclear region (N). Scale bar: 20 μm. In panel **c** and **f**, the boxes show median and 1^st^ and 3^rd^ quartile and the whiskers extending 1.5 times the interquartile range from the boxes. Statistical analysis is performed using One-Way ANOVA followed by Bonferroni post-hoc analysis. **P < 0.01 and ***P < 0.001. Source numerical data are available in Source data.

We next assessed defects in global proteostasis state by monitoring ubiquitin-positive (Ub+) and autophagy receptor p62/SQSTM1 puncta formation. In the fibroblasts, there were no obvious Ub+ or p62/SQSTM1 puncta under basal conditions. When exposed to proteotoxic stress, i.e. sub-lethal dosages of proteasome inhibitor Bortezomib (BTZ) or lysosome inhibitor Chloroquine (CQ), fibroblasts from aged donors showed a moderate increase in an accumulation of Ub+ and p62/SQSTM1 and a much higher increase in those of aged/sAD donors (Extended Data Fig. 1a,b). Our results indicate an expected correlation of fibroblast’s proteostasis vulnerability with donor’s age and disease status, as previously described in human brain^22–24^.

We next generated tNeurons at post-induction day (PID) 35 to 42 (Fig. 1d, and Extended Data Fig. 2a,b). Overall, our protocol can efficiently transdifferentiate fibroblasts, with a slight reduction in fibroblast-to-neuron conversion efficiency in aged donors (Extended Data Fig. 2c,d). tNeurons retained epigenetic hallmarks (Fig. 1e,f). Surprisingly, in aged and AD tNeurons the proteostasis deficits were exhibited under basal conditions, unlike what is observed in the originating fibroblasts. Thus, Ub+ and p62/SQSTM1 puncta were robustly increased in aged/sAD tNeurons (Fig. 1g, and Extended Data Fig. 3a). To assess if aged and AD tNeurons exhibit constitutive deficits in other proteostasis pathways, we monitored the levels of small heat shock protein HspB1 (Extended Data Fig. 3b). There was an increase of HspB1 levels in aged and aged/AD tNeurons as compared to young tNeurons. These experiments indicate that fibroblasts and the derived neurons retain hallmarks of aging and sAD, and that age- and AD-dependent proteostasis deficits become exacerbated in neurons.

### Proteotoxic inclusions formed in aged/sAD tNeurons

We next examined AD-associated protein pathologies in tNeurons. We found dramatic increases in aged/sAD tNeurons for deposits of intra-cellular total Aβ and toxic isoform Aβ42 as well as hyper-phosphorylated Tau (pTau) (Fig. 1g, and Extended Data Fig. 3a). A sensitive ELISA detection assay confirmed increased Aβ42 levels in lysates of aged/sAD tNeurons (Extended Data Fig. 3c). TDP-43 deposits are a pathological hallmark of amyotrophic lateral sclerosis (ALS) and frontotemporal dementia (FTD)^25^, but occur in 23-50% of AD cases^26, 27^. We also observed increased TDP-43 pathology in aged/sAD tNeurons, including cytoplasmic mislocalization of nuclear TDP-43 and hyper-phosphorylated TDP-43 (pTDP-43) (Fig. 1g, and Extended Data Fig. 3a). pTau inclusions partially colocalized with p62/SQSTM1 puncta while pTDP-43 partially colocalized with Ub+ puncta (Extended Data Fig. 3d). Together, these experiments indicate that deficits in neuronal proteostasis are exacerbated by aging and AD, cooperatively promoting the cell-intrinsic accumulation of multiple AD-related protein pathologies.

### Quantitative proteomics of young, aged and aged/sAD tNeurons

Transcriptome of AD patient-derived neurons has been extensively characterized, but often does not reflect the state of their proteome^28–30^. Accordingly, we carried out quantitative proteomic analyses of young, aged and aged/sAD tNeurons. Notably, the top-ranked pathways altered by aging and AD included proteostasis and organelle homeostasis (Fig. 2a, and Extended Data Fig. 4a-c). We observed age-related up-regulation of proteins modulating mitochondria and synapse, while proteins in the endosome-lysosomal pathway (e.g. CLU, CTSC and TMEM175) were mostly down-regulated with aging. Comparing aged vs. young tNeurons and aged/sAD vs. young tNeurons changes revealed shared aged and aged/sAD protein hits, with over 94% of them showing the same direction of expression changes in both aged and aged/sAD tNeurons (Extended Data Fig. 4d,e). Comparing aged/sAD vs. aged proteomic changes revealed sAD-specific changes including up-regulation of proteins annotated as endosome-lysosome (e.g. CLU), mitochondria (e.g. CHDH), inflammation (e.g. PYCARD) and synapse (e.g. SORCS2) as well as sAD-specific down-regulation of membrane and vesicular trafficking proteins (Fig. 2a).

**Fig. 2.**
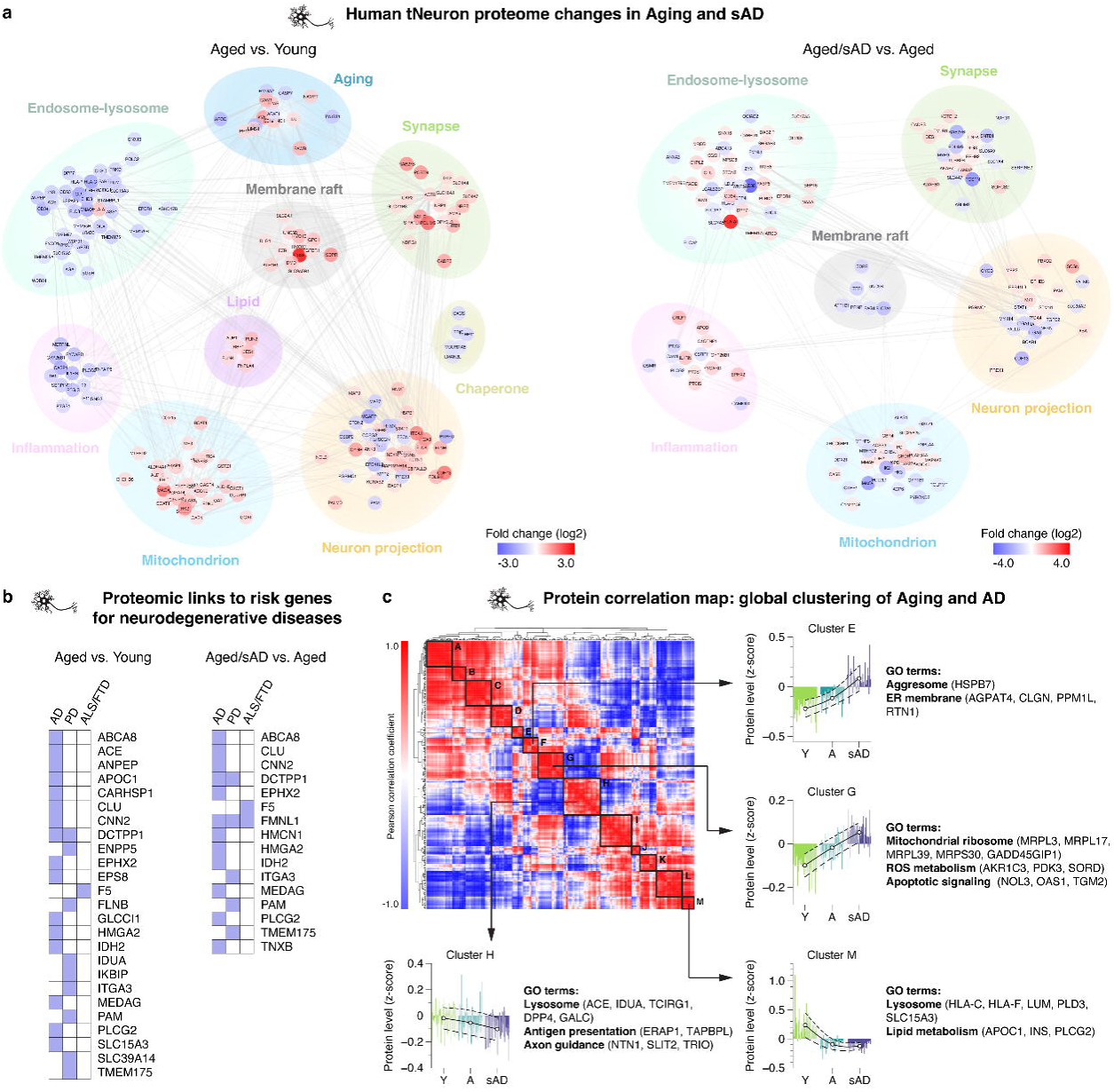
Human tNeurons carry proteomic signatures of aging and AD. **(a)** Differential expression of proteins detected in tNeurons from healthy young (*n* = 3) and aged (*n* = 3) individuals, and patients with aged/sAD (*n* = 6) at PID 40. The top pathways for aging and sAD proteomes are analyzed using gene ontology (GO) databases. Comparison is performed between tNeurons from aged and young donors, and between aged/sAD and aged donors. Colored circles represent the enrichment of identified proteins revealing by log2-fold change: increase in red and decrease in blue. **(b)** List of differentially expressed proteins associated with risk genes for age-related neurodegenerative diseases. AD: Alzheimer’s disease. PD: Parkinson’s disease. ALS/FTD: Amyotrophic lateral sclerosis/Frontotemporal dementia. **(c)** Clustering heatmap of Pearson correlation coefficients of total tNeuron protein expression. Cluster A to M show distinct protein expression patterns and the associated GO terms between young, age and aged/sAD. Each line represents the expression of individual protein defined by the relative protein abundance (z-score) across different groups. White circles represent the average z-score for each cluster. Dash lines represent ±SD.

Remarkably, many aged/sAD proteome changes corresponded to proteins encoded by genes associated with risk of neurodegenerative diseases (e.g. AD, PD and FTD, Fig. 2b), mostly involved in the endosome-lysosomal pathway.

Cluster analysis of our proteomic data based on the similarity of protein expression across young, aged and aged/sAD tNeurons identified related subsets of proteins with unique trajectories of change during aging and AD (Fig. 2c). Clusters E and G exhibited increased protein expression going from young to aged to aged/sAD tNeurons. In contrast, clusters H and M exhibited gradual decreased protein expression. These clusters included proteins regulating lysosome and lipid metabolism. Indeed, the endosome-lysosomal system appears to be a major pathway affected by aging and AD, with many changed proteins associated with lysosomes, notably involved in lysosomal quality control (LQC) altered in aging (e.g. CNN2, HspB1) and AD (e.g. DPP7, PLBD2, PLD3, TAGLN).

### PSEN1 mutations elicit pathological and proteome alterations

We next examined tNeurons derived from fibroblasts of fAD-PSEN1 donors. *PSEN1* mutants elicit early development of AD through increased production of Aβ42^31^. fAD-PSEN1 tNeurons obtained from middle-aged donors also manifested basal accumulation of Ub+ and p62/SQSTM1 and contained comparable or higher levels of Aβ, pTau, p-TDP43 and HspB1 than aged/sAD tNeurons (Extended Data Fig. 5a-c). We next compared the proteomic analysis of fAD-PSEN1 tNeurons with the proteomes of either aged/sAD tNeurons (Extended Data Fig. 5d-f) or young tNeurons (Extended Data Fig. 5g-i) to identify top candidate hits and common top-ranked pathways. Consistent with the younger age of the donors, fAD-PSEN1 tNeurons had reduced expression of proteins in the “Aging” category (Extended Data Fig. 5e). The most dramatic change in fAD-PSEN1 tNeurons was the down-regulation of mitochondrial proteins.

### Constitutive lysosomal damage augmented in aged/sAD tNeurons

Given the changes in the lysosomal proteome in aged and aged/sAD tNeurons, we used transmission electron microscopy (TEM) to characterize lysosomal ultrastructure (Fig. 3a). Comparing young, aged and aged/sAD tNeurons revealed progressive increases in the size of individual lysosomes and increased presence of electron-dense granules. Small electron-dense granules were specifically found proximal to the lysosomal membrane in aged tNeurons, whereas large dense granules accumulated within lysosomes of aged/sAD tNeurons. Notably, aged and aged/sAD tNeurons contained mitochondria closely surrounding the enlarged lysosomes within the cell body (see red arrow in Fig 3a, right panel). Quantification supported an increase in mitochondria-lysosome contacts in AD neurons.

**Fig. 3.**
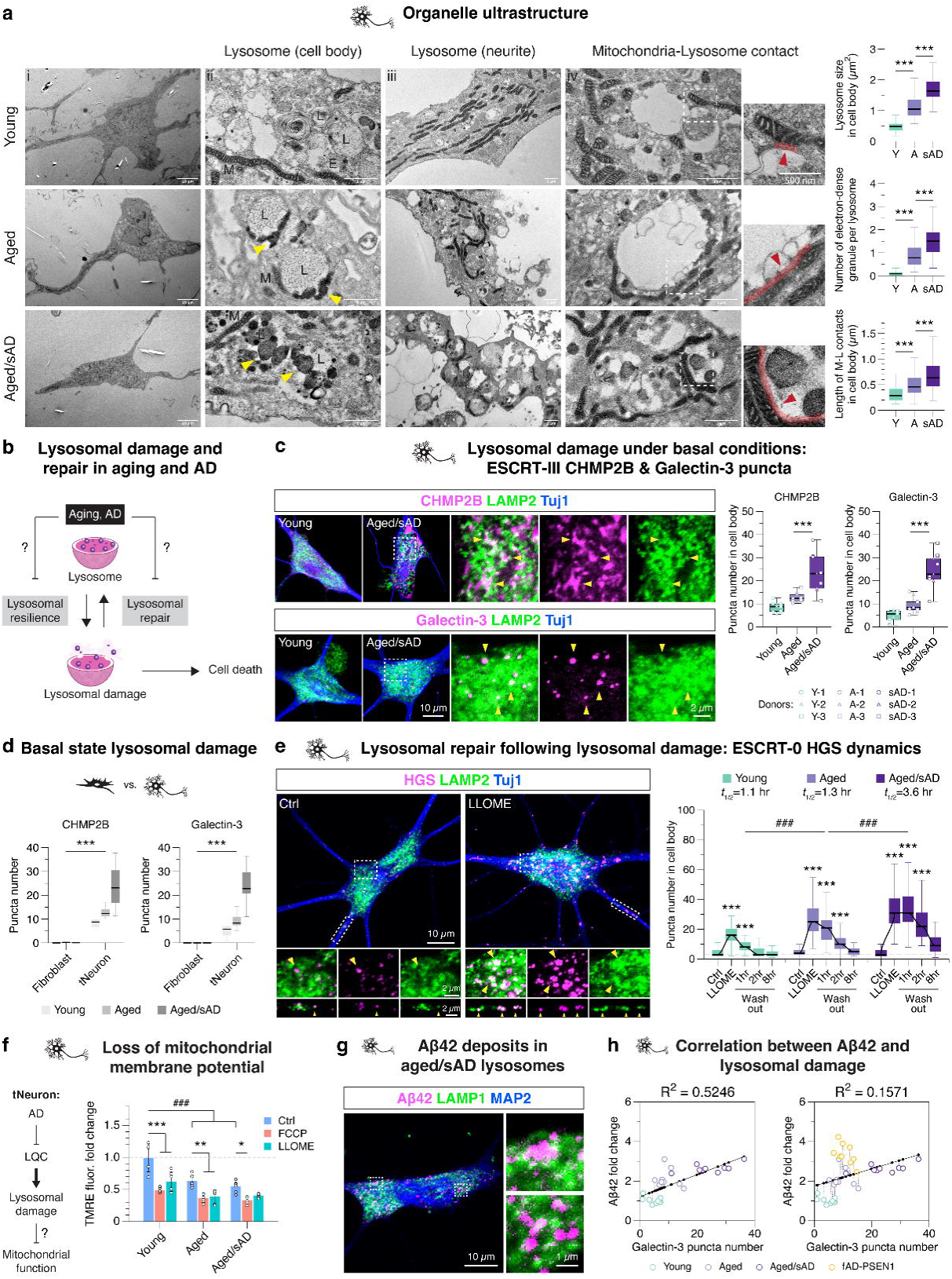
Constitutive lysosomal damage and lysosomal repair deficits in AD tNeurons. **(a)** Transmission electron microscopy (TEM) for analyzing organelle ultrastructure in human tNeurons. E: endosome; L: lysosome; M: mitochondria. Yellow arrowhead: electron-dense granules. Red arrowhead and line: mitochondria-lysosome contact site. Insert: higher magnification view of mitochondria-lysosome contact. Lysosome size: *n* = 59 (young), 69 (aged) and 69 (aged/sAD) lysosomes; electron-dense material abundance: *n* = 74 (young), 68 (aged) and 60 (aged/sAD) lysosomes; mitochondria-lysosome contacts: *n* = 174 (young), 262 (aged) and 246 (aged/sAD) contacts from two donors and two independent experiments. Scale bar (i): 20 μm. Scale bar (ii-iv): 1 μm. **(b)** Testing how aging and AD alters lysosomal damage responses, leading to cell death. **(c)** Representative images of tNeurons immunostained for LAMP2 (green), ESCRT-III CHMP2B and Galectin-3 (magenta) and Tuj1 (blue) at PID 35 at basal conditions (scale bar: 10 μm). IF quantification of numbers of CHMP2B and Galectin-3 puncta in the cell body of tNeurons. CHMP2B: *n* = 117 (young), 103 (aged) and 97 (aged/sAD) cells; Galectin-3: *n* = 111 (young), 108 (aged) and 95 (aged/sAD) cells from three donors and three independent experiments. Insert: higher magnification view of protein colocalization (scale bar: 2 μm). Arrowhead: CHMP2B and Galectin-3 colocalization with LAMP2. **(d)** Comparison of numbers of CHMP2B and Galectin-3 puncta between fibroblasts and tNeurons at basal conditions. Fibroblasts: CHMP2B: *n* = 102 (young), 105 (aged) and 99 (aged/sAD) cells; Galectin-3: *n* = 102 (young), 105 (aged) and 99 (aged/sAD) cells. tNeurons: CHMP2B: *n* = 117 (young), 103 (aged) and 97 (aged/sAD) cells; Galectin-3: *n* = 111 (young), 108 (aged) and 95 (aged/sAD) cells. All data are acquired from three donors and three independent experiments. **(e)** Representative images of AD tNeurons immunostained for LAMP2 (green), ESCRT-0 HGS protein (magenta) and Tuj1 (blue) at PID 36 (scale bar: 10 μm). Cells are treated with L-leucyl-L-leucine O-methyl ester (LLOME) at 0.25 mM for 30 min, and then washed out of LLOME for lysosomal repair. IF quantification of numbers of HGS puncta in the cell body. The half-life (*t_1/2_*) represents the time required for lysosomal repair. Young: *n* = 108 (Ctrl), 107 (LLOME), 111 (Washout 1 hr), 118 (Washout 2 hr) and 90 (Washout 8 hr) cells; Aged: *n* = 110 (Ctrl), 113 (LLOME), 114 (Washout 1 hr), 116 (Washout 2 hr) and 109 (Washout 8 hr) cells; Aged/sAD: *n* = 115 (Ctrl), 115 (LLOME), 117 (Washout 1 hr), 109 (Washout 2 hr) and 79 (Washout 8 hr) cells from three donors and three independent experiments. Insert: higher magnification view of HGS and LAMP2 in the cell body and neurites (scale bar: 2 μm). Arrowhead: HGS colocalization with LAMP2. **(f)** Testing if defective LQC mediates mitochondrial dysfunction in aging and AD. Quantification of mitochondrial membrane potential using TMRE staining after the treatment of DMSO Ctrl, 20 µM FCCP or 0.25 mM LLOME for 30 min. The values are revealed by a fold change relative to young tNeurons treated with DMSO. *n* = 6 (young), 6 (aged) and 6 (aged/sAD) independent replicates from three donors and two experiments. **(g)** IF analysis of colocalization of Aβ42 with LAMP1 in aged/sAD tNeurons (scale bar: 10 μm). Insert: higher magnification view of Aβ42 and LAMP1 (scale bar: 1 μm). **(h)** Analysis of correlation between intra-cellular Aβ42 levels and Galectin-3 puncta numbers in different groups of tNeurons. *n* = 9 independent replicates from three donors and three experiments. Black solid line represents the fitted linear correlation. Coefficient of Discrimination (R^2^) is calculated using Pearson’s correlation. In panel **a**, **c**, **d**, and **e**, the boxes show median and 1^st^ and 3^rd^ quartile and the whiskers extending 1.5 times the interquartile range from the boxes. In panel **f**, data are displayed as mean ± SD. Statistical analysis is performed using One-Way ANOVA (**a**, and **c**) or Two-Way ANOVA (**d, e** and **f**) followed by Bonferroni post-hoc analysis. *P < 0.05, **P < 0.01 and ***P < 0.001. ###P < 0.001. Source numerical data are available in Source data.

Based on our proteomic and TEM analyses, we hypothesized that the cellular state of aged and AD tNeurons affects lysosomal resilience to damage and/or restoration from damage (Fig. 3b). Complex LQC machineries recognize damaged lysosomes to facilitate either repair or clearance and protect cells against lysosomal membrane permeabilization and cell death^32–35^. Two well-characterized mechanisms involve ESCRT proteins (ESCRTs) and Galectins, which target mildly and severely damaged lysosomes, respectively (Extended Data Fig. 6a). To assess lysosomal integrity under basal conditions, we first measured the constitutive recruitment of ESCRTs and Galectins to lysosomes. We detected no appreciable levels of damaged lysosomes in young tNeurons, a slight increase in the number and intensity of ESCRT-III CHMP2B- and Galectin-3-containing puncta in aged tNeurons and a dramatic increase in these lysosomal damage markers in aged/sAD tNeurons (Fig. 3c, and Extended Data Fig. 6b,c). We also found aberrant accumulations of CHMP2B adjacent to the plasma membrane and neurite branch points (Extended Data Fig. 6b), supporting the evidence that CHMP2B participates in both plasma and lysosomal membrane repair^34–36^. Nonetheless, measuring activated Caspase-3/7 levels showed no spontaneously apoptotic death in all tNeuron groups (Extended Data Fig. 6d). These results indicate that aged/sAD tNeurons carry a significant burden of constitutively damaged lysosomes. Interestingly, similar analyses in the parental fibroblasts did not reveal significant CHMP2B- or Galectin-3-positive puncta for any donor groups under basal conditions, in contrast to what we observed in tNeurons (Fig. 3d).

### Aging and sAD impact neuronal lysosomal repair pathways

We next examined if lysosomal repair dynamics are impaired in aged and aged/sAD tNeurons. Thus, constitutive lysosomal damage in aged and aged/sAD tNeurons could overwhelm the LQC machineries and limit cellular capacity to counter additional lysosomal stress. Lysosomal damage was induced by a 30-min incubation with a well-validated lysosomotropic reagent, L-leucyl-L-leucine O-methyl ester (LLOME)^37–39^, followed by a chase for up to 8 hr after LLOME washout to assess repair kinetics. Given the robust baseline presence of ESCRT-III CHMP2B puncta in aged/sAD tNeurons, we chose ESCRT-0 HGS to monitor the spatiotemporal change of lysosomal damage response, because its baseline distribution was comparable across all tNeuron groups (Fig. 3e). Lysosomal damage caused by LLOME treatment indeed increased HGS puncta number; when compared to young tNeurons, aged tNeurons showed a moderate increase and aged/sAD tNeurons showed a substantial increase. Following LLOME washout, the half-life estimates (*t*_1/2_) of HGS puncta decrease revealed differences in the efficiency of lysosomal repair. In young tNeurons, HGS puncta rapidly returned to baseline levels with *t*_1/2_ of ∼1 hr. However, aged tNeurons exhibited a slight delay, whereas aged/sAD tNeurons exhibited a 3-fold delay (Fig. 3e). These experiments indicate that lysosomal repair pathways are progressively impaired in aged and aged/sAD tNeurons.

### Lysosomal damage impacts other proteostasis pathways

Lysosomal damage and repair have been linked to several proteostasis pathways^40–42^ and to RNA-containing stress granules^32^. We thus examined how LLOME-induced damage affects RNA-binding protein TDP-43 and molecular chaperone Hsp70 in the different tNeuron classes. While cytoplasmic TDP-43 was basally increased in aged/sAD tNeurons (Fig. 1g), we observed that TDP-43 mislocalized to damaged lysosomes, particularly in LLOME-treated aged/sAD tNeurons (Extended Data Fig. 6e). Cytoplasmic Hsp70 was basally diffuse in all tNeuron classes and remained diffuse in LLOME-treated young tNeurons but was strongly recruited to damaged lysosomes in aged/sAD tNeurons (Extended Data Fig. 6e). These experiments suggest that lysosomal damage likely reverberates through the network, affecting other proteostasis processes.

Because aged/sAD tNeurons have altered mitochondrial proteomes and increased mitochondria-lysosome contacts, we considered a potential interplay between lysosomal damage and mitochondrial dysfunction in AD. We used tetramethylrhodamine ethyl ester (TMRE), which accumulates only in metabolically active mitochondria. As expected, TMRE fluorescence was reduced in tNeurons treated with an uncoupler of the mitochondrial respiratory chain, FCCP (Fig. 3f). Under basal conditions, TMRE fluorescence was significantly lower in aged and aged/sAD tNeurons compared to young tNeurons, indicative of basal mitochondrial impairment. Lysosomal damage with a 30-min LLOME treatment also led to decreased TMRE fluorescence intensity in all tNeuron classes, revealing a link between lysosomal and mitochondrial impairment.

Finally, we asked if lysosomal damage affects lysosomal acidification in tNeurons. Compared to aged tNeurons, aged/sAD tNeurons exhibited a slight loss of lysosomal acidification at baseline, which was exacerbated by LLOME treatment (Extended Data Fig. 6f). Since previous studies found that APP derivatives preferentially accumulate in poorly acidified lysosomes^43^, we examined if Aβ42 deposits are proximal to lysosomes in aged/sAD tNeurons. Indeed, both APP-CTF and Aβ42 colocalized with LAMP1 (Fig. 3g and Extended Data Fig. 6g) or LC3B (Extended Data Fig. 6h) in aged/sAD tNeurons, supporting a link between lysosomal damage and amyloid accumulation.

### Aging and genetic link to organellar and calcium defects

We next investigated organelle homeostasis in fAD-PSEN1 tNeurons. Unlike aged/sAD tNeurons, fAD-PSEN1 tNeurons contained relatively lower-density accumulation of CHMP2B- and Galectin-3-puncta under basal conditions (Extended Data Fig. 7a). However, the analysis of LLOME pulse-chase lysosomal repair kinetics revealed some similarities between aged/sAD and fAD-PSEN1 tNeurons as both classes formed higher number of HGS puncta than young tNeurons during LLOME treatment (Extended Data Fig.7b). Following LLOME washout, the repair kinetics of both AD tNeuron classes were slow relative to young tNeurons, the *t*_1/2_ of HGS puncta dissipation increased by ∼2-fold in fAD-PSEN1 and ∼3-fold in aged/sAD tNeurons.

Analysis of lysosomal acidification revealed similarly phenotypic parallels between fAD-PSEN1 and aged/sAD tNeurons (Extended Data Fig. 7c). Mitochondrial membrane potential was significantly reduced in both aged/sAD and fAD-PSEN1 tNeurons at basal conditions and declined further after FCCP or LLOME treatment (Extended Data Fig. 7d). While aging seems to be a primary driver of constitutive lysosomal damage, both sAD and fAD-PSEN1 exhibit impaired LQC and reduced mitochondrial metabolism.

Since previous studies revealed lysosomal calcium dysregulation underlying the pathological process in AD^44, 45^, we also measured lysosomal calcium stores in tNeurons using Cal-520, a fluorogenic calcium-sensitive indicator, conjugated with Dextran molecules. Compared to young tNeurons, we found much lower lysosomal calcium stores in aged tNeurons and even less in aged/sAD and fAD-PSEN1 tNeurons at basal conditions (Extended Data Fig. 7e). Since lysosomal acidification and calcium dysregulation are associated with neurons affected by AD, we calculated the correlation between intra-cellular Aβ42 and either lysosomal acidification or lysosomal calcium in young, aged and aged/sAD tNeurons. There was a much stronger correlation between Aβ42 deposits and lysosomal calcium dysfunction than with impaired lysosomal acidification (Extended Data Fig. 7f). Thus, decreased lysosomal calcium stores in tNeurons accompanies the increased vulnerability of the lysosome system in aged and AD tNeurons as well as the increased Aβ42 burden. Nonetheless, fAD-PSEN1 tNeurons had a much higher Aβ42 burden than expected from either lysosomal acidification and calcium deficits from the above correlation. These experiments suggest a nuanced synergy between aging and AD to disrupt organelle homeostasis and lead to cell autonomous proteotoxic Aβ42 deposits.

### Correlation between lysosomal damage and Aβ deposits in tNeurons

The causal relationship between lysosomal damage and Aβ deposition in AD remains an important and poorly understood question^10–12, 43, 46, 47^. Our characterization of both constitutive lysosomal damage and Aβ burden in the same set of tNeurons offered the possibility to assess their correlation in aged and AD tNeurons. When comparing young, aged and aged/sAD tNeurons, we observed a moderate positive correlation between Aβ42 deposits and Galectin-3 (R^2^ = 0.52; Fig. 3h) or CHMP2B puncta number (R^2^ = 0.34, Extended Data Fig. 8a). A similar positive correlation was also observed between these lysosomal damage markers with intra-cellular total Aβ (Extended Data Fig. 8b). As above, the PSEN1-mutant fAD-PSEN1 tNeurons were outliers, as they contained a higher Aβ42 burden than expected from the correlation measured for aged and aged/sAD tNeurons.

We also examined the correlation between lysosomal damage markers CHMP2B and Galectin 3 and deficits in either lysosomal acidification or calcium stores. The comparison of pH-sensing FITC-Dextran fold change showed mild positive correlation with both Galectin-3 or CHMP2B puncta number (Extended Data Fig. 8c). Of note, Calcium-sensing Cal-520 fold change showed a stronger negative correlation with the damage markers (Extended Data Fig. 8d). In all cases, the correlation was ageing- and sAD-dependent. These analyses indicate lysosomal deficits in aging and AD are linked to lysosomal damage.

### Lysosomal damage pathology in AD mouse and patient brains

To link our *in vitro* findings to *in vivo* pathophysiology, we carried out histopathological analyses of post-mortem brain cortex of AD patients and mouse models of aging and AD (Fig. 4a). Compared to young mice (3 month), aged mice (20 to 24 month) exhibited a moderate increase in Galectin-3 immunoreactivity, which colocalized with enlarged LAMP1-positive lysosomes in neurons (Extended Data Fig. 9a,b). We also examined brains from an AD mouse model expressing human APP carrying the Swedish (K670N/M671L) and London (V717I) mutations (APP^Lon/Swe^)^48^. Similar to aged/sAD tNeurons, APP^Lon/Swe^ brains exhibited widespread accumulation of LAMP1-positive clumps colocalizing with CHMP2B, Galectin-3 and Hsp70, which were absent in non-transgenic (NTg) mice brain (Fig. 4b, and Extended Data Fig. 9c-f). These damaged lysosomal clumps were observed intra-cellularly, in the perinuclear region within individual pyramidal neurons as well as in the extra-cellular space devoid of MAP2 staining, suggestive of severe neuronal death linked to lysosomal damage. Of note, pyramidal neurons proximal to Aβ plaques contained intra-neuronal Aβ42 deposits within lysosomes in APP^Lon/Swe^ mice (Fig. 4b).

**Fig. 4.**
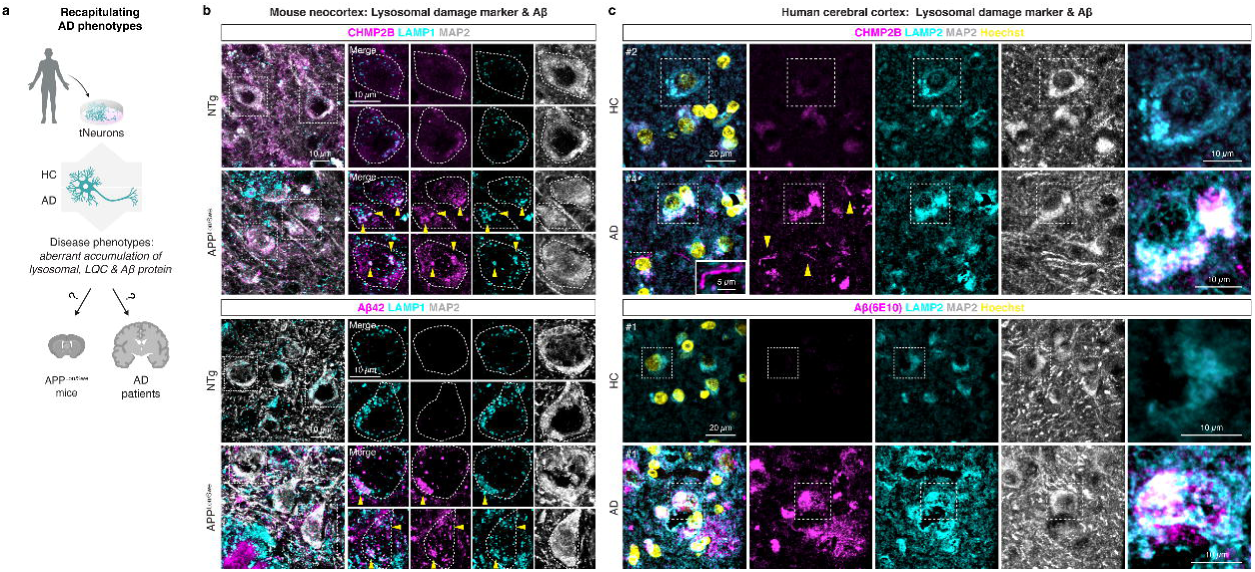
Lysosomal damage is linked to amyloid accumulation in post-mortem brain tissue. **(a)** Schematic for describing an experimental pipeline to test if the disease phenotypes observed in AD tNeurons are also detected in brain tissue of AD patients and transgenic mice expressing mutant human APP with the Swedish (K670N/M671L) and London (V717I) mutations (APP^Lon/Swe^) for modeling AD. HC: healthy control. **(b)** IF staining of CHMP2B, Aβ42 and LAMP1 in the neocortex of non-transgenic mice (NTg) and APP^Lon/Swe^ transgenic mice (scale bar: 10 μm). The brain tissue is co-stained with MAP2. Insert: higher magnification view of colocalization between CHMP2B, Aβ42, LAMP1 within individual neurons (scale bar: 10 μm). Arrowhead: intra-neuronal colocalization of CHPM2B and Aβ42 with LAMP1. **(c)** IF quantification of CHMP2B, Aβ(6E10) and LAMP2 in the cerebral cortex of HC and AD donors (scale bar: 20 μm). The brain tissue is co-stained with MAP2 and Hoechst. Insert: higher magnification view of CHMP2B and Aβ(6E10) colocalization with LAMP2 (scale bar: 10 μm). Arrowhead: CHMP2B-positive fibril structures.

AD patient brain tissue also exhibited intra-neuronal and global elevation of Aβ, CHMP2B and Galectin-3 immunoreactivity colocalized with LAMP2, compared with healthy control (HC) individuals (Fig. 4c, and Extended Data Fig. 10a-d). Similar to our findings in tNeurons, we observed increased colocalization of CHMP2B and Galectin-3 with LAMP2 in AD brains. Interestingly, we identified two distinct patterns of CHMP2B staining in AD patient brains. CHMP2B mainly colocalized with LAMP2-positive lysosomes in the perinuclear region of neurons but also formed thin, thread-like structures of variable length devoid of LAMP2 staining. Using the 6E10 antibody, we observed intra-neuronal and extra-cellular Aβ/APP deposits associated with LAMP2-positive lysosomes in the brains of AD patients (Fig. 4c, and Extended Data Fig. 10c). Despite their complexity, the brains of AD patients and mice models reveal intra-neuronal and global increase in lysosomal damage accompanying co-aggregation with amyloid plaques and neuritic degeneration that are consistent with the phenotypes of our tNeuron models.

### Cell-autonomous inflammatory activation in AD tNeurons

Our proteomic analysis showed that aged/sAD tNeurons upregulate a protein network involved in cytokine signaling, including inflammasome adaptor protein PYCARD/ASC (Fig. 5a). While activation of NLRP3 inflammasomes in microglia has been extensively linked to AD^49^, we asked if this inflammatory response is also elevated in AD neurons and associated with lysosomal damage (Fig. 5b). Immunofluorescence analyses showed that, under basal conditions, aged/sAD donors contained slightly more inflammasome-positive neurons than aged and fAD-PSEN1 donors (Fig. 5c). Following a 3-hr LLOME treatment, the inflammasome-positive neurons were increased by 35-40% in aged/sAD tNeurons compared with 18-30% in aged and fAD-PSEN1 tNeurons. We conclude AD neurons have cell-autonomous inflammasome activation, potentially associated with their higher vulnerability to lysosomal damage.

**Fig. 5.**
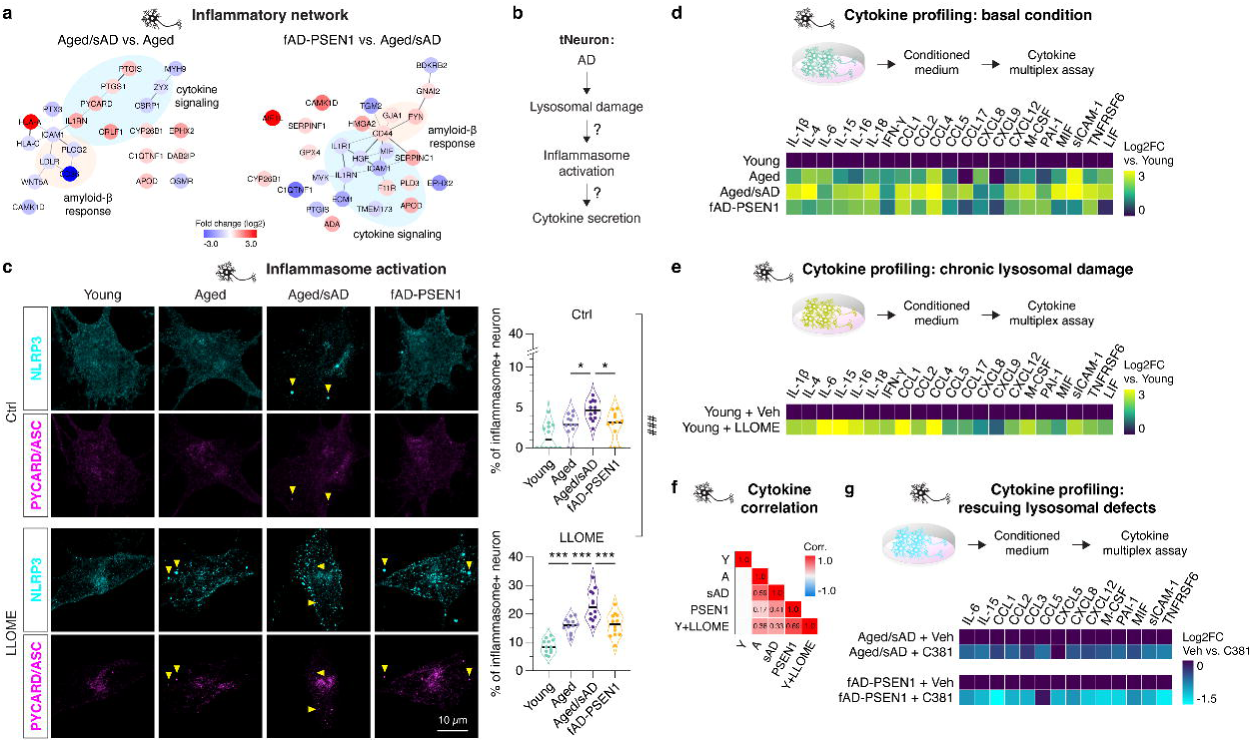
Lysosomal damage mediates inflammatory responses in AD tNeurons. **(a)** Interaction network for proteins involved in the inflammatory response pathway in tNeurons. The relative abundance indicated by log2-fold change (Log2FC): increase in red and decrease in blue. **(b)** Testing if lysosomal damage links to inflammasome activation and cytokine secretion in AD neurons. **(c)** Representative images of tNeurons immunostained for inflammasome markers NLRP3 (cyan) and PYCARD/ASC (magenta) with or without 0.25 mM LLOME treatment for 3 hr at PID 40. IF quantification of the percentage of tNeurons showing inflammasomes per image. *n* = 441 to 477 (young), 399 to 513 (aged) and 522 to 648 (aged/sAD) cells from four donors and three independent experiments. Data are displayed as violin plot indicating median. Arrowhead: colocalization of NLRP3 and PYCARD/ASC. Scale bar: 10 μm. **(d)** Inflammatory profiling of the conditioned medium from all groups of tNeurons at basal conditions at PID 40. Heatmap represents Log2FC in cytokine and chemokine levels relative to young tNeurons. *n* = 6 (young), 6 (aged), 12 (aged/sAD) and 4 (fAD-PSEN1) independent replicates from two experiments. **(e)** Inflammatory profile of the conditioned medium from young tNeurons treated with or without chronic lysosomal damage stress (0.1 mM LLOME starting at PID 33 for 7 days). Log2FC is relative to DMSO (vehicle). *n* = 6 (young + vehicle and 8 (young + LLOME) independent replicates from two experiments. **(f)** Heatmap and Pearson correlation analysis for identified cytokines and chemokines. **(g)** Inflammatory profiling of the conditioned medium from aged/sAD and fAD-PSEN1 tNeurons at PID 35 treated with or without 3.1 µM C381 for 7 days. Log2FC is relative to DMSO (vehicle). *n* = 4 (aged/sAD: vehicle, C381) and 4 (fAD-PSEN1: vehicle, C381) independent replicates from two experiments. Log2-fold change in mean fluorescence intensity (MFI) is used for comparison. Statistical analysis is performed using two-sided Student’s *t*-test (**g**) or One-Way ANOVA (**d**, and **e**) or Two-Way ANOVA (**c**) followed by Bonferroni post-hoc analysis. *P < 0.05 and ***P < 0.001. Source numerical data are available in Source data.

We next examined whether AD tNeurons secrete inflammatory factors. Multiplex cytokine profiling of the conditioned medium from untreated tNeurons using Luminex assays showed that aged/sAD tNeurons have increased secretion of pro-inflammatory (e.g. IL-1β) and anti-inflammatory (e.g. IL-15) cytokines and chemokines (e.g. CCL2) (Fig. 5d, and Supplementary Fig. 1). To assess if lysosomal damage indeed promotes secretion of inflammatory factors, we subjected young tNeurons to low-dose LLOME treatment to elicit a chronic, sublethal lysosomal damage state. This treatment led to increased secretion of IL-1β, IL-6, IFN-γ and CCL2 (Fig. 5e, and Supplementary Fig. 1). Using Pearson’s correlation analysis, we identified a moderate and positive correlation between cytokine secretion, aging, AD and lysosomal damage (Fig. 5f). This led us to hypothesize that rescuing lysosomal deficits should reduce secretion of inflammatory factors. We thus treated tNeurons with C381, a lysosome-targeting small molecule that promotes lysosomal acidification and resilience to damage^50^. Strikingly, the amelioration of lysosomal damage by C381 in in aged/sAD and fAD-PSEN1 tNeurons significantly reduced the secretion of IL-6, IL-15 and CCL2 (Fig. 5g, and Supplementary Fig. 2). These experiments link aging- and AD-dependent lysosomal damage to neuron-autonomous secretion of inflammatory cytokines.

To relate these cell-based findings to physiological disease-relevant context, we examined cytokine expression in published human datasets of single-nucleus transcriptomic analysis in post-mortem cortex from HC donors (Extended Data Fig. 10e). Neurons were confirmed to express cytokine and chemokine transcripts, albeit at lower levels than microglia, which have high transcript levels given their immune-active function^28^. We also conducted an unbiased analysis of CSF proteins in 50 HC and 29 AD donors in a search for biomarkers (Supplementary Table 3). Of note, the elevation of IL-15 was found in both the conditioned medium from aged/sAD tNeurons and CSF from AD patients (Fig. 5d, Extended Data Fig. 10f), has been positively correlated with age of onset in AD^51^.

### Rescuing lysosomal function ameliorates AD pathologies

We next used C381 to further investigate the link between lysosomal impairment, lysosomal dysfunction and neuronal cell death in AD tNeurons (Fig. 6a). Pre-incubation of aged and AD tNeurons with C381 concurrently reduced the number of constitutive CHMP2B and Galectin-3 puncta on lysosomes (Fig. 6b), supporting its ability to rescue lysosomal deficits. We next measured how C381 treatment of tNeurons affects additional lysosomal functions. Since lysosomal hydrolases, such as Cathepsin-B, are optimally active at pH 4-5, lysosomal de-acidification caused by aging, AD or LLOME should abrogate their activity. Indeed, Cathepsin-B hydrolysis of fluorogenic substrate Magic Red was reduced in aged and aged/sAD tNeurons under basal conditions and was dramatically lost upon LLOME treatment (Fig. 6c). However, C381 treatment restored Cathepsin-B activities in LLOME-treated AD tNeurons to near-normal levels. The lysosomal vulnerability of AD tNeurons renders them exquisitely sensitive to LLOME-mediated cell death, as measured by Caspase-3/7 activation (Fig. 6d). This phenotype was abrogated by C381 rescue of lysosomal function, which was strongly neuroprotective from lysosome-mediated apoptosis in AD tNeurons.

**Fig. 6.**
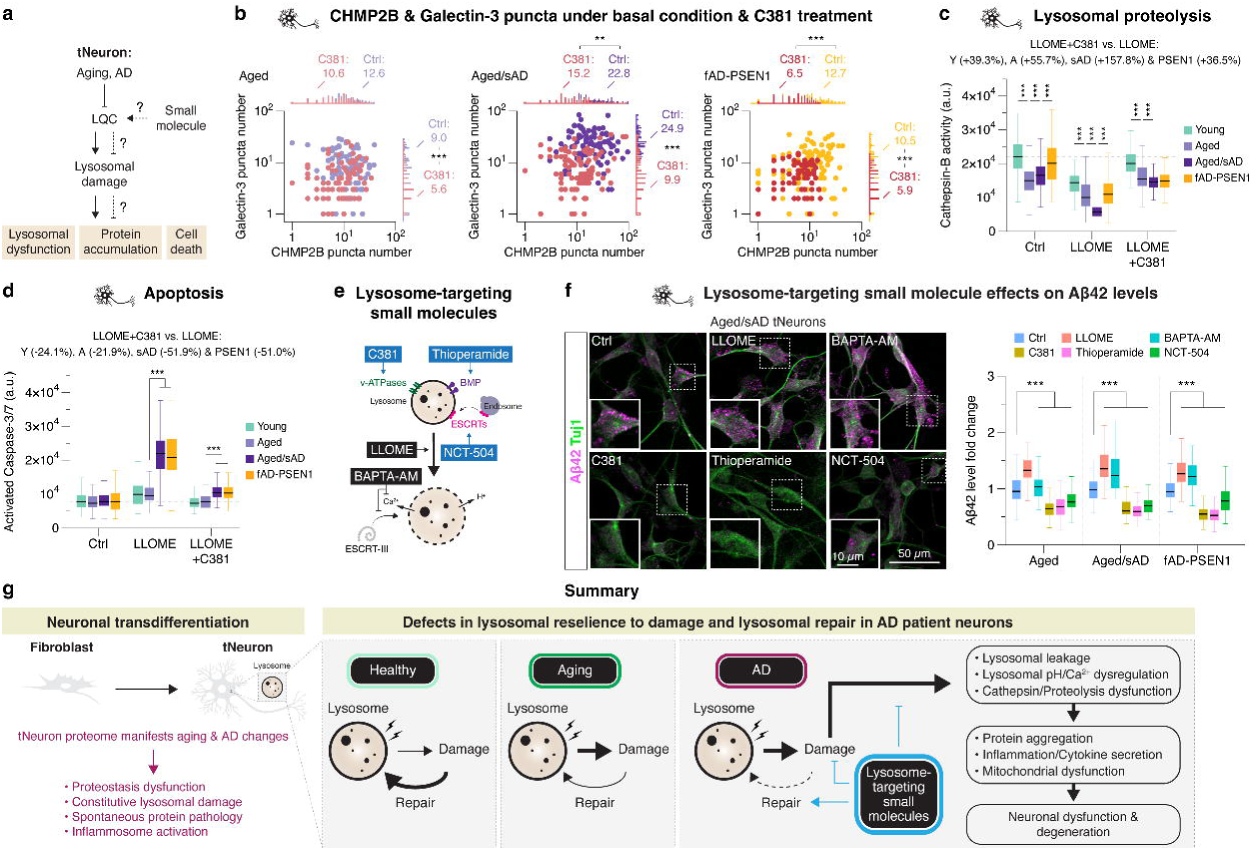
Pharmacological improvement of lysosomal resilience to damage ameliorates AD phenotypes in tNeurons. **(a)** Testing if rescuing LQC provides neuroprotective effects in AD tNeurons. **(b)** Concurrent change of CHMP2B and Galectin-3 puncta number by 0.25 mM LLOME treatment for 30 min at PID 35 following the pre-treatment with DMSO Ctrl or 3.1 µM C381 for 7 days. Ctrl: *n* = 103 (aged), 95 (aged/sAD) and 104 (fAD-PSEN1); C381: *n* = 88 (aged), 95 (aged/sAD) and 91 (fAD-PSEN1) cells from three donors and three independent experiments. Each dot represents the number of detectable CHMP2B (x-axis) and Galectin-3 (y-axis) puncta in individual neuron. **(c)** Measurement of changes in Cathepsin-B activity caused by 0.25 mM LLOME treatment for 30 min at PID 35 following the pre-treatment with 3.1 µM C381 for 7 days. Young: *n* = 145 (Ctrl), 142 (LLOME) and 147 (LLOME+C381) cells; Aged: *n* = 157 (Ctrl), 156 (LLOME) and 122 (LLOME+C381) cells; Aged/sAD: *n* = 157 (Ctrl), 152 (LLOME) and 155 (LLOME+C381) cells; fAD-PSEN1: *n* = 142 (Ctrl), 142 (LLOME) and 141 (LLOME+C381) cells from three donors and three independent experiments. **(d)** Measurement of Caspase-3/7 activation after 0.5 mM LLOME treatment for 1 hr at PID 42. Pre-treatment of 3.1 µM C381 at PID 35 is continued for 7 days. Young: *n* = 162 (Ctrl), 142 (LLOME) and 105 (LLOME+C381) cells; Aged: *n* = 158 (Ctrl), 146 (LLOME) and 133 (LLOME+C381) cells; Aged/sAD: *n* = 148 (Ctrl), 167 (LLOME) and 142 (LLOME+C381) cells; fAD-PSEN1: *n* = 136 (Ctrl), 147 (LLOME) and 148 (LLOME+C381) cells from three donors and three independent experiments. **(e)** Schematic for small molecules that modulate lysosomal function and damage. Small molecules with beneficial effects labelled with blue color, whereas with detrimental effects labelled with black color. **(f)** Effects of small molecules (LLOME: 0.25 mM, BAPTA-AM: 2.5 µM, C381: 3.1 µM, Thioperamide: 5 µM and NCT-504: 2.5 µM for 2-day treatment) on intra-neuronal Aβ42 levels in aged, aged/sAD and fAD-PSEN1 tNeurons. Representative images of aged/sAD tNeurons immunostained for Aβ42 (magenta) and Tuj1 (green) at PID 35 (scale bar: 50 μm). IF quantification of fold-changes in Aβ42 levels during small molecule treatment relative to DMSO Ctrl. *n* = 104 to 122 (aged), 88 to 144 (aged/sAD) and 61 to 136 (fAD-PSEN1) cells from three donors and three independent experiments. Insert: higher magnification view of Aβ42 within individual neuron (scale bar: 10 μm). **(g)** We propose that in AD, either as a result of stochastic events or mutational burdens, lysosomal repair defects are exacerbated, leading to overwhelmed LQC machineries and sustained presence of damaged lysosomes. Restoring lysosomal homeostasis and damage ameliorate AD pathologies in neurons. In panel **c**, **d**, and **f**, the boxes show median and 1^st^ and 3^rd^ quartile and the whiskers extending 1.5 times the interquartile range from the boxes. Statistical analysis is performed using two-sided Student’s *t*-test (**b**) or One-Way ANOVA (**c**, **d**, and **f**) followed by Bonferroni post-hoc analysis. **P < 0.01 and ***P < 0.001. Source numerical data are available in Source data.

The availability of small molecules that can either improve or further impair lysosomal homeostasis was next exploited to test whether lysosomal dysfunction contributes to cell-autonomous Aβ42 deposits in aged and AD tNeurons (Fig. 6e,f). To increase lysosomal damage, we used LLOME or calcium chelator, BAPTA-AM. These treatments significantly increased intra-neuronal Aβ42 levels in both aged/sAD and fAD-PSEN1 tNeurons. To improve lysosomal function, we used three mechanistically distinct compounds. In addition to C381, we used thioperamide, which increases lysosomal phospholipid BMP^52^ and NCT-504, which upregulates ESCRT transcripts^53^. Strikingly, incubating AD tNeurons with either of these compounds significantly reduced the levels of Aβ42 deposits in aged and AD tNeurons by ∼20-46% (Fig. 6f). These experiments support a causal connection between neuronal lysosomal dysfunction and cell-intrinsic generation of intra-neuronal Aβ42 deposits. They further demonstrate that ameliorating lysosomal function can by itself reduce Aβ42 burdens in AD neurons.

## Discussion

Here we demonstrate that neuronal transdifferentiation provides a powerful approach to study cellular and mechanistic aspects of human brain aging and AD. While iPS cells have great potential to model human genetic disorders^54^, the cellular rejuvenation process erases access to the contribution of aging to neurodegenerative diseases^15, 16, 55, 56^. Building on previous studies showing that human tNeurons preserve hallmarks of aging^16, 56–58^, we harnessed tNeurons to reveal proteomic signatures and identify proteostasis and LQC pathways as selectively affected by aging and AD (Fig. 6g). Our proteomic analyses resonate with the GWAS and PWAS studies identifying AD causal and risk genes belonging to the endosome-lysosomal pathway, such as *CLU*, *PLD3* and *SNX32*^8, 59^. We propose that LQC defects are an early pathogenic event that causes neurons to experience sustained stress and deterioration, leading to cell-autonomous formation of proteotoxic deposits, mitochondrial dysfunction, inflammasome activation and risk of neurodegeneration. This may suggest that aging and AD-linked mutations act in a two-hit process. In the first hit, neurons enter a vulnerable phase with impaired proteostasis and lysosomal homeostasis. Persistent lysosomal damage and LQC defects diminish neuronal resilience to counteract harmful insults, such as mutations or sporadic stressful events, that would constitute the second hit, triggering the degenerative process. In turn, these defects may then elicit cell non-autonomous mechanisms, such as inflammation or spreading of toxic aggregates, that aggravate overt neuronal loss across brain regions. Our work further suggests that tNeurons could be a reliable tool for evaluating small molecules possessing neuroprotective effects against AD pathologies.

Age-dependent impairment of lysosomal function is observed along a longitudinal aging axis in different model systems^3, 5, 60^. Lysosomal function is also impaired in AD, and brain tissues from AD patients are extensively characterized by the co-occurrence of Aβ and lysosomal pathology including amyloid plaques enriched in lysosomal proteins^61–63^. The causal relationship between amyloid accumulation and lysosomal damage is still poorly understood. There is a long-standing hypothesis that increased endocytosis of extra-cellular protein oligomers causes lysosomal damage, leading to seeding of protein aggregates and ultimately neuronal death in AD^10–12^. Recent studies reported that intrinsically perforated endosome-lysosomes are present in diseased neurons and facilitate the seeding of cytoplasmic aggregates following internalization of pre-formed fibrils^64, 65^. Our tNeuron models provide valuable insights into this important problem.

Here, we show that constitutive disruptions in lysosomal membrane integrity and reduced lysosomal repair mechanisms are increased by aging and more by late-onset sAD. These phenotypes are correlated with, but may not solely depend on intra-cellular Aβ. One possibility is that the tNeuron phenotypes reflect a very early stage of dysfunction in AD. While we do not seed with fibrils in this study, it is possible that lysosomal dysfunction leads to elevated Aβ levels that are then secreted from AD tNeurons; subsequent uptake of extra-cellular Aβ would further aggravate lysosomal damage phenotypes. The vicious cycle proposed by our model would eventually lead to severe collapse of endosome-lysosomal homeostasis essential for neuronal survival. Importantly, our experiments also suggest this vicious cycle might be interrupted by small molecules that ameliorate lysosomal function.

The measurement of *in vivo* lysosomal pH in transgenic mouse models for AD indicated that the emergence of lysosomal acidification deficits precedes Aβ depositions and that as disease progresses, neurons build up Aβ/APP-CTF selectively in perinuclear de-acidified lysosomes, leading to lysosomal damage^43^. However, our tNeuron models do not fully recapitulate this phenotype. We only observed a subtle deficit in lysosomal acidification in AD tNeurons at baseline, similar to previous findings in cells lacking *PSEN1/2* or *PLD3*, gene variants linked to early- and late-onset AD, respectively^66, 67^ but we detect a greater lysosomal acidification deficit under stressful conditions. This could be because tNeurons detect early manifestations in AD pathology and investigating early triggers of endosome-lysosomal dysfunctions present significant challenges in transgenic AD mice or human brains. LQC machineries are likely severely impaired by the time of analysis, rendering them incapable of restoring lysosomes, resulting in significant lysosomal de-acidification. Interestingly, we observed lysosomal calcium dysregulation in aged and AD tNeurons at basal conditions and found it better correlated with lysosomal damage and intra-cellular Aβ levels. This raises an important point remaining to be determined concerning the relative contribution of lysosomal acidification or calcium deficits as early triggers of AD^67, 68^. There is an increasing attention on intra-cellular accumulation of amyloids in AD, associated with intrinsic lysosomal defects at early disease stages^43, 46^. Mass cytometry analysis of human post-mortem brain found that neurons accumulating intra-cellular Aβ are preferentially lost early during the progression of AD in contrast to tau, that preferentially accumulates in neuronal sub-types resilient to neuronal loss^46^. The accumulation of Aβ/APP-CTF promotes neuropathy by eliciting endosomal abnormality and Rab5 overactivation and compromising lysosomal calcium stores through inhibition of lysosome-ER contacts in *PSEN1* knockout or mutant mouse neurons or human iNeurons^45, 69–71^. Interestingly, we found that the severity of lysosomal phenotypes in fAD-PSEN1 tNeurons much resemble those of aged tNeurons, suggesting that this genetic variant and aging make comparable contribution to neuronal lysosomal defects, at least under basal conditions. When exposed to additional insults, aged/sAD and fAD-PSEN1 tNeurons are undoubtfully highly vulnerable to lysosomal stress. Since PSEN1 is directly involved in APP processing and Aβ generation, it is possible that in these cells Aβ accumulation precedes lysosomal dysfunction, and these deposits eventually overwhelm LQC and lead to constitutive lysosome damage.

Damage accumulation is an inevitable feature of aging in living organisms. To ensure cellular homeostasis, it is essential to repair and remove damaged proteins and/or organelles to slow aging. Growing evidence indicate that lysosomal clearance declines with age and age-related diseases^72, 73^. However, it is yet to be understood how LQC pathways are affected by aging and human disease. Our results uncover an age-dependent decline in ESCRT- and Galectin-mediated LQC that promotes cell-autonomous proteostasis deficits, cytokine secretion and cell death in tNeurons of AD patients. Of note, recent studies suggested that molecular chaperones, PI4K2A, ORP family members, LRRK2, Annexin A7 and stress granules, are also involved in LQC^32, 33, 74–77^. Due to the increasing complexity of the LQC machinery, it would be of interest to evaluate other LQC pathways to understand their impact on aging and AD in the future. Surprisingly, our findings that CHMP2B, LAMP1, Hsp70, Aβ, and TDP-43 become associated with damaged lysosomes in neurons are reminiscent of phenotypes linked to granulovacuolar degeneration (GVD). GVD is one of typical AD hallmarks, characterized by the presence of cytoplasmic granule-containing vacuoles, termed GVD bodies^78^. Revealed by neuropathological observations in post-mortem brain tissue of AD patients, GVD bodies are selective to pyramidal neurons, structurally resemble endocytic and autophagic vesicles and contain a variety of proteins, including the typical hallmark proteins as aforementioned^78, 79^. However, the pathological mechanisms of their formation remain mostly unclear. Our *in vitro* and *in vivo* findings may suggest that lysosomal damage could be involved in GVD etiology during the development of AD. In addition, our tNeuron proteomics revealed a decrease in levels of PLD3 in aging and sAD. PLD3 is a lysosomal exonuclease enriched in the senile plaques in the brain of AD patients^66, 80, 81^. The levels of PLD3 transcripts and proteins are reduced in the brain of AD patients. Depletion of PLD3 led to lysosomal dysfunction and lipid accumulation, and activation of the cGAS–STING signaling mediated by cytosolic nucleotides leaked out from lysosomes^66^. STING is a substrate of lysosomal degradation to attenuate the activation of cGAS–STING signaling and control pro-inflammatory cytokine expression^82^. However, STING is also a driver of autophagy activation. The failure in lysosomal degradation of STING aberrantly activates Atg5-dependent autophagy, which amplifies immune responses and a variety of cell death pathways^83^. Previous findings also indicated that lysosomal membrane permeabilization in microglia activates NLRP3 inflammasomes and promotes the secretion of pro-inflammatory cytokines, thus aggravating neuronal damage^84^. Our study indicates that constitutive lysosomal damage promotes inflammasome activation in AD tNeurons, consistent with a recent study that indicates AD tNeurons recapitulate senescence-like neurons observed in AD patient brain as a novel source of neuroinflammation^20^.

There are limitations to our study. We use monocultures of tNeurons without glial cell support and short-term culture conditions (i.e. 5 to 6 weeks), which are quite distinct from the complex tissue organization and prolonged aging processes in human brain. Thus, our transdifferentiated cell-based models may not entirely reflect the physiological signatures of aging and AD. The neurons are derived from human adult fibroblasts of several donors to account for patient heterogeneity. The challenging conversion and cell culture system limit the experimental scalability for larger donor cohort sizes and large-scale biochemical analyses. In addition, healthy young and aged fibroblasts were collected from donors who are clinically normal at collection, but we cannot rule out their possibility of developing dementia and AD due to unidentified genetic variations or environmental factors in subsequent decades. Aging affects many branches of proteostasis^5, 85, 86^, as demonstrated by our finding of increased p62, Ub+ and HspB1 with aging and AD, as well as increases in pathogenic protein deposits and impaired lysosomal and mitochondria function. Therefore, cellular health relies on the interplay among different proteostasis pathways, raising the question of which are the upstream triggers of aging-associated proteostasis decline. How the entire LQC pathways are impaired by aging and AD remains to be comprehensively investigated in future studies.

In sum, we propose that tNeurons provide insights into early neuron-intrinsic cell biological processes by which loss of proteostasis and organelle homeostasis contribute to AD pathogenesis. These insights would be impossible to obtain in intact brains, where the complex interplay between cell types in the tissue establishes a vicious cycle that likely exacerbates all responses. They would also not be possible in stem cell-derived neurons, which lack the aging-linked phenotypes essential to these late-onset diseases, such as AD. One corollary of our experiments is that counteracting intrinsic proteostasis and lysosomal homeostasis deficits in aged and AD tNeurons may be attractive strategies for early-stage prevention of the cascade of deleterious events in affected AD brains.

## Acknowledgments

We thank the Stanford Alzheimer’s Disease Research Center (ADRC), D. Channappa and M. Greicius with kind support for collecting and identifying human fibroblast lines, post-mortem brain tissue and CSF samples. Y. Rosenberg-Hasson and J. Liang from Stanford Human Immune Monitoring Center for help with cytokine profiling; J. Perrino from Stanford Cell Sciences Imaging Core Facility for help with TEM experiments; P. Chu from Stanford Human Pathology/Histology Service Center for help with tissue processing; M. Weglarz from Stanford Shared FACS Facility for FACS support. We thank T.K. Rainbolt, V.B. Masto and S. Taguwa of the Frydman laboratory for helpful discussions on experimental design; R.I. Morimoto, S. Finkbeiner and D. Gate for helpful inputs to the manuscript. Figure illustrations in Fig. 1a, 3b, 4a and Extended Data Fig. 1b, 9a,c were generated in part with BioRender (https://www.biorender.com/). This work was supported by NIA P01AG054407 (J.F., D.F. & J.W.K.); the Hevolution Foundation (J.F., M.A.P., D.F. & J.W.K.); the Michael J. Fox Foundation for Parkinson’s Research and the Aligning Science Across Parkinson’s (ASAP) initiative (J.F.). The Michael J Fox Foundation administers the grant ASAP-000282 on behalf of ASAP and itself; the Glenn Foundation for Medical Research Postdoctoral Fellowship in Aging Research (C.-C.C.); the Paul F. Glenn Center for Biology of Aging Research at Stanford (T.W.-C. and JF); the Life Sciences Research Foundation Postdoctoral Fellowship (C.-C.C.) and the Alzheimer’s Disease Research program in BrightFocus Foundation Grant (C.-C.C.); the NIA-funded Stanford ADRC P50AG047366 and P30AG066515 (T.W.-C. & P.M.L.); the Goldberg Fund Fellowship and the Miguel Servet Program (CP21/00017) (M.A.P.).

## Author contributions

C.-C.C. and J.F. conceptualized the study, designed experiments, interpreted data and wrote the manuscript with inputs from all authors. C.-C.C. developed tNeurons, conducted cell culture, immunofluorescence staining, microscopy, flow cytometry, conditioned medium experiments with the help of T.-T.L.. C.-C.C. performed transdifferentiation with the help of Y.S. under the supervision of M.W.. C.-C.C., R.V. and J.L. carried out mice and drug experiments under the supervision of T.W.-C.. M.A.P. and J.W.-G. set up the proteomic data-processing pipelines and performed bioinformatic data analysis with the help of C.-C.C. and J.A.P. under the supervision of S.P.G. and D.F.. P.M.L. processed and analyzed human CSF data under the supervision of T.W.-C.. J.W.K. provided key reagents and helped design lysosome function assays. All authors read and approved the manuscript.

## Methods

Experiments involving cell culture studies were conducted according to a protocol reviewed and approved by Stanford University. All animal care and procedures complied with the Animal Welfare Act and were in accordance with institutional guidelines and approved by the V.A. Palo Alto Committee on Animal Research and the institutional administrative panel of laboratory animal care at Stanford University. Details of key resources used in this study are provided in a Key Resource Table available on Zenodo: 10.5281/zenodo.14606908. Detailed protocols used in this study can be accessed on protocols.io as listed below: dx.doi.org/10.17504/protocols.io.36wgq3edklk5/v1, dx.doi.org/10.17504/protocols.io.kxygx317wg8j/v1, dx.doi.org/10.17504/protocols.io.j8nlko6d5v5r/v1, dx.doi.org/10.17504/protocols.io.dm6gp356pvzp/v1, dx.doi.org/10.17504/protocols.io.n2bvj3qm5lk5/v1.

### Experimental Model and Subject Details

#### Human subjects

De-identified human fibroblasts, post-mortem prefrontal cortex and CSF samples were acquired from individuals of various ages and disease conditions from the Stanford Alzheimer’s Disease Research Center (ADRC). Cells and tissue samples were obtained under written consents from all subjects approved by Institutional Review Board of Stanford University. The cell and tissue samples collected by ADRC were not specifically for this study. For histological experiments, of these subjects, 8 were assessed as healthy control (HC) and 10 were patients with cognitive impairment (dementia due to AD). For CSF proteomics experiments, 50 were assessed as HC and 29 were patients with cognitive impairment (MCI or dementia due to AD). Age and sex demographics are detailed in Supplementary Table 1 and 3. In the Stanford ADRC and Coriell Institute, fibroblasts were collected from cognitive normal young and aged donors, and individuals with sAD or fAD showing the clinical symptoms of AD, including progressive cognitive impairment. In the ADRC, individuals with cognitive decline received neurological examinations and cognitive tests to determine cognitive status and consensus diagnosis by a team of neuropathologists. The pathological diagnosis of post-mortem tissues was made by microscopic examination of multiple brain regions using Amyloid score, Braak neurofibrillary degeneration score and CERAD neuritic plaque score.

#### HEK293T cells

The HEK293T cells are the cell line derived human embryonic kidney and were acquired directly from ATCC (CRL-1573). Cells were grown in culture medium (DMEM supplemented with GlutaMAX, 10% FBS, 1% Penicillin-Streptomycin, 1% HEPES and 1% Sodium pyruvate) (Thermo Fisher Scientific) sterilized by a 0.22 µm vacuum filter (Thermo Fisher Scientific) in a 37°C incubator with 5% CO_2_ in the air. The HEK293T cells were used for lentiviral production by transfecting lentiviral vector of interest mixed with packaging and envelope plasmid. Passaging the cells was performed with Trypsin-EDTA (Thermo Fisher Scientific) every three days.

#### Human fibroblasts

Primary human adult fibroblasts derived from clinically healthy adults and individuals diagnosed with AD were collected from shared resources in the Stanford ADRC and Coriell Institute for Medical Research, which operates the NIGMS, NIA, NINDS cell repository. Culture of primary human fibroblasts was described at https://dx.doi.org/10.17504/protocols.io.36wgq3edklk5/v1. Briefly, cells were grown in culture medium (DMEM supplemented with GlutaMAX, 10% FBS, 1% Penicillin-Streptomycin, 1% MEM NEAA, 1% Sodium pyruvate and 0.1% β-mercaptoethanol (BME)) (Thermo Fisher Scientific) sterilized by a 0.22 µm vacuum filter and maintained in a 37°C, 5% CO_2_ incubator. The subculture of proliferating fibroblasts used for regular experiments and neuronal transdifferentiation was typically within 3 to 7 passages. Some fibroblast lines obtained with slightly higher passage numbers were used for neuronal transdifferentiation no more than 12 passages. Passaging the cells was performed with Trypsin-EDTA every 6 to 7 days.

#### Human tNeurons

tNeurons were converted from human fibroblasts with low passage numbers using a combination of transcription factors and small molecules. This method was derived from the protocols as previously described^17, 87^ and modified for this study. A detailed protocol can be found at https://dx.doi.org/10.17504/protocols.io.j8nlko6d5v5r/v1. Briefly, the lentiviral expression of *Brn2*, *Ascl1*, *Myt1l* and *Ngn2* (also referring as BAMN factors) in human fibroblasts initiated neuronal reprogramming. Transduced cells underwent puromycin and PSA-NCAM selection and were cultured in reprogramming medium (DMEM/F12:Neurobasal (1:1) medium, 2% B-27, 1% N-2, 0.25% GlutaMAX, 1% Penicillin-Streptomycin) (Thermo Fisher Scientific) for 15 days and then switched to maturation medium (BrainPhys neuronal medium (STEMCELL Technologies), 2% B-27, 1% N-2, 0.25% GlutaMAX, 1% Penicillin-Streptomycin) for additional 15 to 22 days. The culture media were supplemented with BDNF and NT-3 (neurotrophic factors; Peprotech), Doxycycline (effector for the Tet-On system; Cayman), Forskolin (cAMP activator; Sigma-Aldrich), SB 431542 (TGF-β/Activin/Nodal inhibitor; TOCRIS), Dorsomorphin (BMP inhibitor; TOCRIS) and XAV939 (Wnt inhibitor; Stemgent; removal from maturation medium). Human tNeurons resemble cortical glutamatergic neurons and were used for the evaluation of reprogramming efficiency, phenotypic characterization, and small molecule treatments after 5 weeks in culture in a 37°C, 5% CO_2_ incubator. The medium was half-changed every 2 to 4 days throughout the lifetime of the culture. More details were described in Method Details > Direct generation of neurons from human fibroblasts.

#### Mice

All mice used in this study were C57BL/6 genetic background. Mice of old age (20 to 24-month old) were obtained from the National Institute on Aging rodent colony and young age (3-month old) were obtained from Jackson Laboratories or Charles River Laboratories. All experiments used male mice. A transgenic mouse model with the expression of high levels of human APP751 carrying V717I, K670M/N671L mutations (also referring to APP^Lon/Swe^) in neurons under control of a Thy1.2 promoter has been studied in different laboratories. The APP^Lon/Swe^ mice developed amyloid plagues associated with an overproduction of Aβ42 in the neocortex and working memory deficits at 3 months old and the plaque formation spread to the hippocampus and thalamus region at 5 to 7 months old^48, 88^. This study used 3 to 6-month-old APP^Lon/Swe^ and the age-matched non-transgenic mice. All mice were kept in a 12-hr light/dark cycle in a temperature- and humidity-controlled environment and provided ad libitum access to food and water.

### Lentivirus preparation

Preparation of lentiviruses was previously described at https://dx.doi.org/10.17504/protocols.io.kxygx317wg8j/v1. The FUW lentiviral vector expressing BAMN factors and EGFP is under the control of TetO promoter and M2rtTA under the control of ubiquitin promoter. HEK293T cells were plated at a density of 6 × 10^6^ in a poly-L-ornithine-coated 10-cm dish and the next day co-transfected with 5 µg lentiviral transfer vector, 4 µg packaging plasmid (psPAX2) and 2.5 µg envelope plasmid (pMD2.G) using Lipofectamine 2000 in OptiMEM (Thermo Fisher Scientific). After 6-hr incubation of Lipofectamine/DNA mixture in OptiMEM, the transfection medium was replaced with fresh DMEM supplemented with GlutaMAX, 2% FBS, 1% Penicillin-Streptomycin, 1% HEPES and 0.1% BME. The cell supernatants containing lentiviral particles were harvested after 24 hr and stored at 4°C. Cells were replenished with fresh DMEM medium with 2% FBS and cultured for additional 24 hr. The supernatants were then harvested and pooled with the first collection. To remove cell debris, the supernatants were centrifugated at 400 x G for 5 min and passed through 0.45 µm syringe filters. The clear virus-containing media can be stored at 4°C for about 1 week. For long-term storage, alternatively, the virus-containing media were spun by ultracentrifugation at 25,000 rpm for 90 min at 4°C to pellet the viruses. The viruses were resuspended in DMEM medium with 2% FBS and snap-frozen in small aliquots to store at −80°C.

### Direct generation of neurons from human fibroblasts

Human adult fibroblasts were plated at a density of 200 × 10^3^ per well of a 6-well plate coated with poly-L-ornithine. The next day, Day 0, fibroblasts were infected with lentiviruses expressing BAMN factors and M2rtTA with or without EGFP by incubating with the diluted virus-containing medium in DMEM supplemented with GlutaMAX, 2% FBS, 1% Penicillin-Streptomycin, 1% HEPES and 0.1% BME plus 4 µg/mL Polybrene for 24 hr. On Day 1, the virus-containing medium was then discarded and replaced by fresh fibroblast culture medium plus 1 µg/mL Doxycycline. On Day 2, puromycin (0.5 μg/mL) was added for selection for 48 hr. On Day 4, the transduced cells were subjected to PSA-NCAM+ selection following the manufacturer’s instructions. Briefly, 0.05% Trypsin-EDTA was added to the cells for 5 min at 37°C to dissociate them from surface and neutralized by fibroblasts culture medium, followed by the centrifugation at 300 x G for 5 min at room temperature to pellet cells. The cells were resuspended in autoMACS buffer and labeled with anti-PSA-NCAM-APC (Miltenyi Biotec) for 10 min at 4°C in the dark. After a wash by autoMACS buffer and centrifugation at 300 x G for 10 min, the cells were incubated with anti-APC MicroBeads (Miltenyi Biotec) for 15 min at 4°C in the dark. Unbound beads were then washed off and the cells were resuspended in autoMACS buffer for subsequent flow cytometry analysis and separation of magnetically PSA-NCAM-labeled and unlabeled cells. The PSA-NCAM+ cells were re-plated at a density of 50 × 10^3^ cells per cm^2^ to the plate coated with vitronectin (VTN-N, 5 μg/mL; Thermo Fisher Scientific) and laminin (rhLaminin-521, 1 μg/mL; Corning). Cells were cultured in fibroblast culture medium plus 1 µg/mL Doxycycline and the next day switched to reprogramming medium (DMEM/F12:Neurobasal (1:1) Medium, 2% B-27, 1% N-2, 0.25% GlutaMAX, 1% Penicillin-Streptomycin) supplemented with small molecules: 1 µg/mL Doxycycline, 5 µM Forskolin, 10 µM SB 431542, 2 µM Dorsomorphin and 2 µM XAV939). After one week, 10 ng/mL BDNF and NT-3 were added to the reprogramming medium. Half of the medium was changed every 2 to 3 days. After 8 days, cells were switched to maturation medium (BrainPhys Neuronal Medium, 2% B-27, 1% N-2, 0.25% GlutaMAX, 1% Penicillin-Streptomycin) supplemented with 1 µg/mL Doxycycline, 5 µM Forskolin, 10 µM SB 431542, 2 µM Dorsomorphin and 10 ng/mL BDNF and NT-3 and cultured for additional 15 to 22 days. Half of the medium was changed every 3 to 4 days. The efficiency of transdifferentiation of human fibroblasts into tNeurons was measured by the percentage of remaining transduced cells that express Tuj1, NeuN and MAP2 and the percentage of EGFP-positive cells showing neuron-like morphology.

### Quantitative proteomics (TMT, Tandem Mass Tag)

Flash frozen cell pellets were lysed in 8M urea buffer (8M urea, 150 mM NaCl, 50 mM HEPES pH 7.5, 1x EDTA-free protease inhibitor cocktail (Roche), 1x PhosSTOP phosphatase inhibitor cocktail (Roche)). Lysates were clarified by centrifugation at 17,000 x G for 15 min at 4°C. Protein concentration of the supernatant was quantified by bicinchoninic acid assay (BCA, Thermo Fisher Scientific). To reduce and alkylate cysteines, 100 µg of protein was sequentially incubated with 5 mM TCEP for 30 min, 14 mM iodoacetamide for 30 min, and 10 mM DTT for 15 min. All reactions were performed at room temperature. Next, proteins were chloroform-methanol precipitated and the pellet resuspended in 200 mM EPPS pH 8.5. Then, LysC (Wako) was added at 1:100 (LysC:protein) ratio and incubated overnight at room temperature in an orbital shaker at 1,500 rpm. The day after, samples were further digested for 5 hr at 37°C with trypsin at 1:75 (trypsin:protein) ratio in an orbital shaker at 1,500 rpm. After digestion, samples were clarified by centrifugation at 17,000 x G for 10 min. Peptide concentration of the supernatant was quantified using a quantitative colorimetric peptide assay (Thermo Fisher Scientific). For TMT labelling, peptides from tNeuron samples were labelled with TMTpro-16plex tags. Methods were previously described for TMT labeling^89–91^. Briefly, 25 μg of peptides was brought to 1 μg/μL with 200 mM EPPS (pH 8.5), acetonitrile (ACN) was added to a final concentration of 30% followed by the addition of 50 μg of each TMT reagents. After 1 hr of incubation at room temperature, the reaction was stopped by the addition of 0.3% hydroxylamine (Sigma) for 15 min at room temperature. Extra information regarding both TMT sample labels is included in Supplementary Table 4. After labelling, all samples were combined, desalted with tC18 SepPak solid-phase extraction cartridges (Waters), and dried in the SpeedVac. Next, desalted peptides were resuspended in 5% ACN, 10 mM NH4HCO3 pH 8 and fractionated in a basic pH reversed phase chromatography using a HPLC equipped with a 3.5 µm Zorbax 300 Extended-C18 column (Agilent). Fractions were collected in a 96-well plate, then combined into 24 samples. Twelve of them were desalted following the C18 Stop and Go Extraction Tip (STAGE-Tip)^92^ and dried down in a SpeedVac. Finally, peptides were resuspended in 1% formic acid, 3% ACN, and analyzed by LC-MS3 in an Orbitrap Fusion Lumos (Thermo Fisher Scientific) equipped with FAIMS and running in RTS-MS3 mode^93–95^. More information regarding all MS parameters is included in Supplementary Table 4. A suite of in-house pipeline (GFY-Core Version 3.8, Harvard University) was used to obtain final protein quantifications from all RAW files collected. RAW data were converted to mzXML format using a modified version of RawFileReader (5.0.7) and searched using the search engine Sequest or Comet^96–98^ against a human target-decoy protein database (downloaded from UniProt in June 2019) that included the most common contaminants. Precursor ion tolerance was set at 20 ppm and product ion tolerance at 1 Da. Cysteine carbamidomethylation (+57.0215 Da) and TMT tag (+304.2071 Da for TMTpro-16pex) on lysine residues and peptide N-termini were set as static modifications. Up to 2 variable methionine oxidations (+15.9949 Da) and 2 miss cleavages were allowed in the searches. Peptide-spectrum matches (PSMs) were adjusted to a 1% FDR with a linear discriminant analysis^99^ and proteins were further collapsed to a final protein-level FDR of 1%. TMT quantitative values we obtained from MS3 scans. Only those with a signal-to-noise ratio > 100 and an isolation specificity > 0.7 were used for quantification. Each TMT was normalized to the total signal in each column. Quantifications included in Supplementary Table 4 are represented as relative abundances. RAW files will be made available upon request. The data have been deposited in the ProteomeXchange Consortium via the PRIDE^100^ partner repository with the data set identifier PXD059089. TMT-proteomics revealed a total of 6,015 proteins with more than two unique peptides, allowing us to identify the top proteomic hits affected by aging and sAD. Biological pathway and gene ontology enrichment analysis were performed using the ClueGo (Cytoscape plug-in), Enrichr or STRING.

### Caspase-3/7 activation

To detect apoptosis in tNeurons, we incubated cells with the Caspase-3/7 substrate FAM-DEVD-FMK (ImmunoChemistry), one of the fluorochrome-labeled inhibitors of caspases that covalently and irreversibly binds to the active caspases. The green fluorescence intensity is a direct measurement of Caspase-3/7 activity. Human tNeurons were treated with DMSO or 0.5 mM LLOME for 1 hr to measure the lysosome-mediated apoptosis or pretreated with 3.1 µM C381 followed by LLOME treatment to assess the rescuing effect of C381 on lysosome-mediated apoptosis. The FAM-DEVD-FMK reagent was reconstituted in DMSO and stored at - 20°C. When cells were available, the FAM-DEVD-FMK reagent was diluted with PBS 1:5 ratio and added to the cell culture medium at a dilution of 1:30 to form 1X staining solution. Cells were incubated with the FAM-DEVD-FMK solution for 30 min at 37°C. After a rinse by Apoptosis Buffer, the cells were fixed by 4% paraformaldehyde (PFA) for 15 min and counterstained with 5 μg/mL Hoechst for confocal microscope imaging.

### Fluorescence-conjugated Dextran assay for measuring lysosomal acidification

Fluorescein isothiocyanate (FITC)-conjugated Dextran at 40 kDa (Thermo Fisher Scientific) were reconstituted in H_2_O and stored at −20°C. Human tNeurons were seeded at 5 × 10^4^ per well of a 24-well plate or 5 × 10^3^ per well of a 96-well plate. Cells were incubated with FITC-Dextran at 0.5 mg/mL for 4 hr at 37°C and rinsed by PBS, followed by a 20-hr chase in fresh culture medium to accumulate Dextran in late endosomes and lysosomes. Then, the cells were treated with DMSO or 0.25 mM LLOME for 30 min. The cells were fixed by 4% PFA for 15 min and prepared for imaging by the confocal microscope and CLARIOstar plate reader.

### Magic Red Cathepsin-B assay for measuring lysosomal proteolysis

Magic Red Cathepsin-B substrate, MR-(RR)2, containing arginine-arginine (RR) sequence was reconstituted in DMSO and stored at −20°C. When cells were available for an experiment, MR- (RR)2 was diluted with H_2_O 1:10 ratio and added to cell culture medium at a dilution of 1:25 to form 1X staining solution. Active Cathepsin-B cleaves MR-(RR)2 and emits fluorescence with optimal excitation of 592 nm and emission of 628 nm. To test lysosomal proteolytic capacity, human tNeurons were incubated MR-(RR)2 staining solution with for 30 min at 37°C, followed by DMSO or 0.25 mM LLOME treatment for 30 min. To evaluate the pharmacological rescuing effect, cells were pre-treated with 3.1 µM C381 for 7 days before the cells were loaded with MR- (RR)2. Cells were treated with 0.25 mM LLOME for 30 min, and then fixed by 4% PFA for 15 min for imaging by the confocal microscope and CLARIOstar plate reader.

### Mitochondrial membrane potential

To detect mitochondrial membrane potential in human tNeurons, we used the tetramethylrhodamine ethyl ester (TMRE) reagent (Abcam), which accumulates in functional and polarized mitochondria according to Δψm. The TMRE reagent was reconstituted in DMSO for a stock solution at 1 mM and stored at −20°C. When tNeurons were in culture for 5 weeks, the cells were pre-treated with DMSO, 50 μM FCCP or 0.25 mM LLOME for 10 min. Then TMRE reagent was added to fresh cell culture medium at a dilution of 1:1000 along with FCCP or LLOME. Half of the old culture medium was replaced with the TMRE-containing medium in order to incubate the cells with TMRE at a final concentration of 500 nM for 30 min at 37°C. Cells were then rinsed with pre-warmed 0.2% bovine serum albumin (BSA)/PBS twice and positioned in the CLARIOstar plate reader for fluorescence measurements with setting of optimal acquisition parameters (excitation of 549 nm and emission of 575 nm).

### ELISA

To measure Aβ42 levels, human tNeurons were trypsinized, washed with ice-cold PBS and pelleted by centrifugation for 5 min at 1,000 X G. Cells were then lysed with RIPA buffer (25 mM Tris-HCl pH 7.5, 150 mM NaCl, 1% NP-40, 0.5% sodium deoxycholate) supplemented with protease inhibitors (Roche). Protein concentrations were determined by BCA assay (Thermo Fisher Scientific). For Aβ42 assay, we used human Aβ42 ELISA Kit (Thermo Fisher Scientific) to detect and quantify the levels in total tNeuron lysates. Briefly, 50 µL of the cell lysates were added to each well of a 96-well plate, followed by the incubation with Aβ42 antibody for 3 hr, anti-rabbit IgG HRP for 30 min and stabilized chromogen for 30 min. The plate was analyzed according to manufacturer’s protocol and Aβ42 values were normalized to total protein concentration of lysates. Two independent experiments and cells from two HC and AD patients with three technical replicates (wells) were performed in this experiment.

### Calcium imaging

Cal-520-Dextran Conjugates at 3 kDa (AAT Bioquest) is a calcium-sensitive dye for detecting intra-cellular calcium levels, particularly in the compartmentalized organelles. Cal-520 was reconstituted in DMSO, aliquoted into single-use volumes and stored at −20°C. Cells were plated on sterile multi-chamber glass bottom slides for 5 weeks and then incubated with 5 μM Cal-520 and 0.1 μM LysoTracker Red DND-99 (Thermo Fisher Scientific) for 2 hr at 37°C. After washing off excess dye by PBS, cells were prepared for live-cell imaging by Zeiss LSM 980 microscope in a 37°C incubation chamber with 5% CO_2_. Cal-520: excitation of 490 nm and emission of 525 nm; LysoTracker Red DND-99: excitation of 577 nm and emission of 590 nm.

### Mice brain perfusion and tissue processing

Mice were anaesthetized with 2.5% (v/v) Avertin (Sigma-Aldrich). Transcardial perfusion with 50 mL cold PBS was performed using a peristaltic pump with the perfusate flow rate not exceeding 10 mL/min. Brain tissue processing was performed as described previously^50, 101^. Hemibrains were isolated and fixed in 4% PFA overnight at 4°C before transferring to 30% sucrose in PBS at 4°C for preservation. Hemibrains were cryosectioned coronally at a thickness of 40 μm on a freezing-sliding microtome, and the free-floating sections were stored in cryoprotectant (40% PBS, 30% glycerol, 30% ethylene glycol) and kept at −20°C until staining.

### Immunofluorescence and image acquisition and analysis

Human adult fibroblasts were fixed with 4% PFA for 15 min at room temperature, permeabilized with 0.3% Triton X-100 in PBS for 5 min or cold Methanol on ice for 20 min, blocked with pre-warmed 5% BSA in PBS and shook for 45 min at room temperature. Fibroblasts were incubated with primary antibody solution in 5% BSA overnight at 4°C in a moisture chamber: anti-CHMP2B (1:400; Proteintech, 12527-1-AP), anti-Galectin-3 (1:1000; BioLegend, 125401), anti-γ-H2AX (1:200; Millipore Sigma, 05-636), anti-H3K9me3 (1:500; Abcam, ab8898), anti-H4K16ac (1:300; Thermo Fisher Scientific, MA5-27794), anti-LAMP1 (1:2000; Cell Signaling, 9091), anti-LAMP2 (1:200; DSHB, H4B4), anti-p62/SQSTM1 (1:400; Abcam, ab56416), anti-S100A4 (1:200; Abcam, ab124805), anti-Ubiquitin (1:200; LifeSensors, AB120), and anti-Vimentin (1:400; Cell Signaling, 5741). Fibroblasts were washed with PBS three times and incubated with fluorophore-conjugated secondary antibody solution in the dark for 1 hr at room temperature: anti-mouse, anti-rabbit and anti-rat (1:500, Thermo Fisher Scientific). Finally, fibroblasts were washed with PBS three times, mounted with DAPI containing ProLong Glass Antifade Mountant (Thermo Fisher Scientific) and air-dried for overnight prior to imaging.

Human tNeurons were undergone two-step fixation: 1) removed half of the culture medium and added equivalent to half the volume of 4% PFA for 5 min at room temperature; 2) removed the old solution and added 4% PFA for 15 min at room temperature. The tNeurons were permeabilized with 0.3% Triton X-100 in PBS for 5 min at room temperature or cold Methanol on ice for 20 min, blocked with pre-warmed 5% BSA in PBS and shook for 45 min at room temperature. tNeurons were incubated with primary antibody solution in 5% BSA overnight at 4°C in a moisture chamber: anti-ASC/PYCARD (1:100; Santa Cruz, sc-514414), anti-GAP-43 (1:300; Novus Biologicals, NB300-143), anti-MAP2 (1:1000; BioLegend, 822501), anti-NeuN (1:500; Abcam, ab177487), anti-NLRP3 (1:200; R&D systems, MAB7578), anti-p62/SQSTM1 (1:500; Proteintech, 18420-1-AP), anti-Amyloid-β (1:100; Cell Signaling Technology, 8243), anti-Amyloid-β (1-42) (1:200; Enzo Life Sciences, ADI-905-804-100), anti-APP-CTF (1:200; BioLegend, 802803), anti-HGS (1:200; GeneTex, GTX101718), <u>anti-Hsp27 (HspB1) (1:100; Proteintech, 18284-1-AP)</u>, <u>anti-Hsp70 (1:1000; Abcam, ab45133),</u> anti-LC3B (1:100; Cell Signaling, 2775), anti-TDP-43 (1:1000; Proteintech, 12892-1-AP), anti-pTau (AT8) (1:100; Thermo Fisher Scientific, MN1020), anti-pTau (S262) (1:200; FUJIFILM WAKO, 010-27123), anti-pTDP-43 Ser409/410 (1:300; Cosmo Bio, CAC-TIP-PTD-M01 & 1:200; BioLegend, 829901), anti-Synapsin-1 (Syn-1) (1:200; Abcam, ab64581), anti-Tau (1:100; Aves Labs), and anti-Beta tubulin III (Tuj1) (1:500; Neuromics, CH23005 & 1:1000; BioLegend, 801201). The tNeurons were washed with PBS three times and incubated with fluorophore-conjugated secondary antibody solution in the dark for 1 hr at room temperature: anti-mouse, anti-rabbit, anti-rat, and anti-chicken (1:500, Thermo Fisher Scientific), washed with PBS three times, mounted with DAPI containing ProLong Glass Antifade Mountant (Thermo Fisher Scientific) and air-dried for overnight prior to imaging.

A detailed protocol for immunostaining of free-floating frozen and paraffin-embedded tissue sections can be found at https://dx.doi.org/10.17504/protocols.io.dm6gp356pvzp/v1. Briefly, free-floating mouse tissue sections at 30 to 50 µm were collected and proceed for immunostaining in multi-well plates. Mouse tissue sections were first rinsed with PBS three times and permeabilized with 0.3% Triton X-100 in PBS for 20 min at room temperature, followed by incubation with blocking buffer (10% Normal Donkey Serum and 0.03% Triton-X-100 in PBS) and shook for 1 hr at room temperature. After PBS rinsing, the tissue sections were incubated with primary antibody solution in 10% Normal Donkey Serum in PBS overnight at 4°C in a moisture chamber: anti-Amyloid-β (1-42) (1:100; Enzo Life Sciences, ADI-905-804-100), anti-CHMP2B (1:100; Proteintech, 12527-1-AP), anti-Galectin-3 (1:50; R&D systems, AF1197), anti-Hsp70 (1:300; Abcam, ab45133), anti-LAMP1 (1:50; Santa Cruz, sc-19992), and anti-MAP2 (1:500; BioLegend, 822501). Fluorophore-conjugated secondary antibody solution was incubated for 1 hr at room temperature: anti-mouse, anti-rabbit, anti-goat and anti-chicken (1:500, Thermo Fisher Scientific). Tissue sections were washed with PBS two times, counterstained with Hoechst (1:2000) and washed with PBS two times again before mounting with ProLong Glass Antifade Mountant (Thermo Fisher Scientific). Samples were air-dried for overnight prior to imaging.

Paraffin-embedded human brain tissues of the cerebral cortex were sectioned at 10 μm thickness. Deparaffinization was achieved by washing slides through xylenes twice, each for 5 min and rehydrating via gradient ethanol (100% ® 95% ® 70% ® 50%) into water, each for 10 min. Following heat-mediated antigen retrieval using Citrate buffer, pH 6.0 (Sigma-Aldrich) for 30 min at 95° C, tissue sections were rinsed with PBS once and incubated with blocking buffer (10% Normal Donkey Serum and 0.03% Triton-X-100 in PBS) for 2 hr at room temperature. Slides were incubated with antibody cocktails in blocking buffer overnight at 4°C in a moisture chamber: Amyloid-β (6E10) (1:100; BioLegend, 803001), anti-CHMP2B (1:100; GeneTex, GTX118181), anti-Galectin-3 (1:50; R&D systems, AF1197 & 1:100; BioLegend, 125401), anti-Hsp70 (1:300; Abcam, ab45133), anti-LAMP2 (1:200; Abcam, ab213294), and anti-MAP2 (1:500; BioLegend, 822501). Tissue sections were incubated with fluorophore-conjugated secondary antibody solution for 1 hr at room temperature: anti-mouse, anti-rabbit, anti-rat, anti-goat and anti-chicken (1:500, Thermo Fisher Scientific). After a PBS rinse, slides were counterstained with Hoechst (1:2000) and washed with PBS two times. The complete air-dried tissue sections were mounted with ProLong Glass Antifade Mountant (Thermo Fisher Scientific) and subjected to dry in the dark overnight prior to imaging.

Imaging was acquired at Z-series (10 to 30 sections; 0.2 to 1 μm steps) according to experimental paradigm using Zeiss LSM 700 and 980 confocal fluorescence microscope with 20x, 63x and 100x objectives. In each experiment, all groups were subjected to image using the same acquisition settings. The z-stack images were performed Maximum Intensity Projection to analyze the mean pixel intensity and determine a threshold to quantify puncta number in the cells by Fiji. For quantitative histology, three to five separate sections were sampled using a 20x objective and fluorescence signals were measured from entire image field to the mean fluorescence change. Tissue sections were imaged and analyzed by blinded observers.

### Transmission electron microscopy

Cells were grown on Ibidi dishes: µ-Dish 35 mm, high Grid-50 Glass Bottom is a 35 mm then fixed in Karnovsky’s fixative: 2% Glutaraldehyde (EMS Cat# 16000) and 4% PFA (EMS Cat# 15700) in 0.1 M Sodium Cacodylate (EMS Cat# 12300) pH 7.4 for 1 hr, chilled and sent to Stanford’s CSIF on ice. They were then post-fixed in cold 1% Osmium tetroxide (EMS Cat# 19100) in water and allowed to warm for 2 hr in a hood, washed 3X with ultra-filtered water, then en bloc stained 2 hr in 1% Uranyl Acetate at room temperature. Samples were then dehydrated in a series of ethanol washes for 10 min each at room temperature beginning at 30%, 50%, 70%, 95%, changed to 100% 2X, then Propylene Oxide (PO) for 10 min. Samples were infiltrated with EMbed-812 resin (EMS Cat#14120) mixed 1:1, and 2:1 with PO for 2 hr each. The samples were then placed into EMbed-812 for 2 hr opened then placed into flat molds w/labels and fresh resin and placed into 65°C oven overnight. Cells of interest were located using the grid pattern and cut out with a gem saw and remounted on pre-labeled resin blocks with fresh resin and polymerized overnight again. Once full polymerized the glass coverslip was etched away using hydrofluoric acid for 20 min. Using the finder grid pattern left behind the block faces were trimmed down allowing for serial sectioning of the cells of interest. Sections were taken around 90 nm, picked up on formvar/Carbon coated slot Cu grids, stained for 40seconds in 3.5% Uranyl Acetate in 50% Acetone followed by staining in 0.2% Lead Citrate for 6 min. Observed in the JEOL JEM-1400 120kV and photos were taken using a Gatan Orius 2k X 2k digital camera.

### Cytokine profiling analysis on neuronal conditioned medium using Luminex multiplex analysis

Secretion of inflammatory factors was analyzed using cytokine profiling of the conditioned medium from tNeurons of healthy donors and AD patients as previously described (https://dx.doi.org/10.17504/protocols.io.n2bvj3qm5lk5/v1). Conditioned medium was collected 48 hr after the last medium change in a 12-well plate with 1 mL of neuronal maturation medium at PID 38, and centrifuged at 10,000 x G for 10 min at room temperature to pellet out particulates. For Human 80 plex panel (EMD-Millipore), a minimum of 200 µL of supernatants was stored at −80°C. Cell free medium was also collected to monitor the background fluorescence. Cell numbers were determined by an automated cell counter for normalization of cytokine levels. The setup of cytokine profiling assay was performed according to the manufacturer’s instructions. Briefly, samples were mixed with antibody-linked magnetic beads on a 96-well plate and incubated overnight at 4°C with shaking. Cold and room temperature incubation steps were performed on an orbital shaker at 500 to 600 rpm. Plates were washed twice with wash buffer in a Biotek ELx405 washer. Following one hr incubation at room temperature with biotinylated detection antibody, streptavidin-PE was added for 30 min with shaking. Plates were washed as above and PBS added to wells for reading in the Luminex FlexMap3D Instrument with a lower bound of 50 beads per sample per cytokine. Each sample was measured in duplicate. Custom Assay Chex control beads were purchased from Radix Biosolutions, Georgetown, Texas, and are added to all wells. The analyses of all conditioned medium samples were performed using raw data (mean fluorescence intensity (MFI)) rather than concentration (pg/mL) to avoid calculating bias per as per recommendation of the Stanford Human Immune Monitoring Center.

### CSF samples and protein discovery

We used the SOMAScan assay platform^102, 103^ (SomaLogic Inc.) to measure the relative levels of 76 human proteins in CSF. This platform is based on modified single-stranded DNA aptamers (SOMAmer) capable of binding to specific protein targets with high sensitivity and specificity. We collected 79 CSF samples (50 HC and 29 AD samples) from a multi-ethnic cohort of older American adults (age range: 60 to 87 years) between 2015 and 2020. Samples were stored at - 80°C and 150 µL aliquots of CSF were sent on dry ice to SomaLogic. CSF samples were analyzed via SOMAScan assay in five batches. To account for variation within and across batches, control, calibrator and buffer samples are added in each 96-well plate. Data normalization was conducted by the manufacturer following three stages. First, in Hybridization Control Normalization, hybridization control probes are used to remove individual sample variance. Second, Intraplate Median Signal Normalization, median normalization removed inter-sample differences within the plate. Last, Plate Scaling and Calibration, this final step removed variance across assay runs.

### Statistics and reproducibility

Quantification of fluorescence images was performed by CLARIOstar plate reader software and open-source Fiji software. For each technical replicate, the fluorescence intensity of the background from cell-free solution or cell-free area in the image field was subtracted from intensity measurements. No statistical method was used to pre-determine the sample size. <u>Representative images shown in</u> Fig. 1d,g, 3g, 4b,c, and Extended Data Fig. 2b, 3d, 6g,h, 9e,f were repeated at least two times. For quantifications of cytokine levels and human and mouse brain samples, the data analyses were blinded to the group allocations. Statistical significance was determined by <u>two-sided Student’s</u> *<u>t</u>*<u>-test</u>, One-Way ANOVA or Two-Way ANOVA based on the experimental design using GraphPad Prism Software. All values were expressed as the box-and-whisker plots or mean ± SD. Differences between two groups were analyzed using <u>two-sided Student’s *t*-test</u> with Welch’s correction. Differences between multiple groups were analyzed using One-Way or Two-Way ANOVA followed by Bonferroni post-hoc analysis. Differences were considered statistically significant for P values < 0.05.

## Data availability

All proteomic source data of human tNeurons have been deposited and are publicly available at ProteomeXchange (accession number: PXD059089). The mass spectrometry parameters, sample information, raw data and the comparison between our datasets and public genomic, transcriptomic and proteomic repositories are provided in Supplementary Table 2 and 4. The cell lines, reagents, plasmids and software presented in the manuscript are reported in Supplementary Table 5 to 9. Source data have been provided in Source Data. The key resource table, supplementary datasets and source data related to this publication are also available from Zenodo.org: 10.5281/zenodo.14606908. All other data supporting the findings of this study are available from the corresponding author on reasonable request. Further requests for resources and reagents should be directed to the Lead Contact, Judith Frydman.

## References (Main)

1. Balch, W.E., Morimoto, R.I., Dillin, A. & Kelly, J.W. Adapting proteostasis for disease intervention. Science (New York, N.Y.) 319, 916–919 (2008).

2. Kaushik, S. & Cuervo, A.M. Proteostasis and aging. Nature medicine 21, 1406–1415 (2015).

3. Hughes, A.L. & Gottschling, D.E. An early age increase in vacuolar pH limits mitochondrial function and lifespan in yeast. Nature 492, 261–265 (2012).

4. Gottschling, D.E. & Nyström, T. The Upsides and Downsides of Organelle Interconnectivity. Cell 169, 24–34 (2017).

5. Lapierre, L.R., Kumsta, C., Sandri, M., Ballabio, A. & Hansen, M. Transcriptional and epigenetic regulation of autophagy in aging. Autophagy 11, 867–880 (2015).

6. Cannizzo, E.S. et al. Age-related oxidative stress compromises endosomal proteostasis. Cell reports 2, 136–149 (2012).

7. Douglas, P.M. & Dillin, A. Protein homeostasis and aging in neurodegeneration. The Journal of cell biology 190, 719–729 (2010).

8. Van Acker, Z.P., Bretou, M. & Annaert, W. Endo-lysosomal dysregulations and late-onset Alzheimer’s disease: impact of genetic risk factors. Molecular neurodegeneration 14, 20 (2019).

9. Aman, Y. et al. Autophagy in healthy aging and disease. Nat Aging 1, 634–650 (2021).

10. Ditaranto, K., Tekirian, T.L. & Yang, A.J. Lysosomal membrane damage in soluble Abeta-mediated cell death in Alzheimer’s disease. Neurobiology of disease 8, 19–31 (2001).

11. Kosenko, E., Poghosyan, A. & Kaminsky, Y. Subcellular compartmentalization of proteolytic enzymes in brain regions and the effects of chronic β-amyloid treatment. Brain research 1369, 184–193 (2011).

12. Zaretsky, D.V., Zaretskaia, M.V. & Molkov, Y.I. Membrane channel hypothesis of lysosomal permeabilization by beta-amyloid. Neuroscience letters 770, 136338 (2022).

13. Calafate, S., Flavin, W., Verstreken, P. & Moechars, D. Loss of Bin1 Promotes the Propagation of Tau Pathology. Cell reports 17, 931–940 (2016).

14. Laskowska, E., Kuczynska-Wisnik, D. & Lipinska, B. Proteomic analysis of protein homeostasis and aggregation. J Proteomics 198, 98–112 (2019).

15. Studer, L., Vera, E. & Cornacchia, D. Programming and Reprogramming Cellular Age in the Era of Induced Pluripotency. Cell Stem Cell 16, 591–600 (2015).

16. Mertens, J. et al. Directly Reprogrammed Human Neurons Retain Aging-Associated Transcriptomic Signatures and Reveal Age-Related Nucleocytoplasmic Defects. Cell Stem Cell 17, 705–718 (2015).

17. Pang, Z.P. et al. Induction of human neuronal cells by defined transcription factors. Nature 476, 220–223 (2011).

18. Mertens, J. et al. Age-dependent instability of mature neuronal fate in induced neurons from Alzheimer’s patients. Cell Stem Cell 28, 1533–1548.e1536 (2021).

19. Victor, M.B. et al. Striatal neurons directly converted from Huntington’s disease patient fibroblasts recapitulate age-associated disease phenotypes. Nature neuroscience 21, 341–352 (2018).

20. Herdy, J.R. et al. Increased post-mitotic senescence in aged human neurons is a pathological feature of Alzheimer’s disease. Cell Stem Cell 29, 1637–1652.e1636 (2022).

21. Capano, L.S. et al. Recapitulation of endogenous 4R tau expression and formation of insoluble tau in directly reprogrammed human neurons. Cell Stem Cell 29, 918–932 e918 (2022).

22. Nativio, R. et al. Dysregulation of the epigenetic landscape of normal aging in Alzheimer’s disease. Nature neuroscience 21, 497–505 (2018).

23. Mansuroglu, Z. et al. Loss of Tau protein affects the structure, transcription and repair of neuronal pericentromeric heterochromatin. Scientific reports 6, 33047 (2016).

24. Salminen, A. et al. Emerging role of p62/sequestosome-1 in the pathogenesis of Alzheimer’s disease. Progress in neurobiology 96, 87–95 (2012).

25. Chou, C.C. et al. TDP-43 pathology disrupts nuclear pore complexes and nucleocytoplasmic transport in ALS/FTD. Nature neuroscience 21, 228–239 (2018).

26. Amador-Ortiz, C. et al. TDP-43 immunoreactivity in hippocampal sclerosis and Alzheimer’s disease. Annals of neurology 61, 435–445 (2007).

27. Josephs, K.A. et al. Staging TDP-43 pathology in Alzheimer’s disease. Acta neuropathologica 127, 441–450 (2014).

28. Grubman, A. et al. A single-cell atlas of entorhinal cortex from individuals with Alzheimer’s disease reveals cell-type-specific gene expression regulation. Nature neuroscience 22, 2087–2097 (2019).

29. Mathys, H. et al. Single-cell transcriptomic analysis of Alzheimer’s disease. Nature 570, 332–337 (2019).

30. Raj, T. et al. Integrative transcriptome analyses of the aging brain implicate altered splicing in Alzheimer’s disease susceptibility. Nature genetics 50, 1584–1592 (2018).

31. Citron, M. et al. Mutant presenilins of Alzheimer’s disease increase production of 42-residue amyloid beta-protein in both transfected cells and transgenic mice. Nature medicine 3, 67–72 (1997).

32. Bussi, C. et al. Stress granules plug and stabilize damaged endolysosomal membranes. Nature 623, 1062–1069 (2023).

33. Tan, J.X. & Finkel, T. A phosphoinositide signalling pathway mediates rapid lysosomal repair. Nature 609, 815–821 (2022).

34. Radulovic, M. et al. ESCRT-mediated lysosome repair precedes lysophagy and promotes cell survival. The EMBO journal (2018).

35. Skowyra, M.L., Schlesinger, P.H., Naismith, T.V. & Hanson, P.I. Triggered recruitment of ESCRT machinery promotes endolysosomal repair. Science (New York, N.Y.) 360 (2018).

36. Jimenez, A.J. et al. ESCRT machinery is required for plasma membrane repair. Science (New York, N.Y.) 343, 1247136 (2014).

37. Thiele, D.L. & Lipsky, P.E. Mechanism of L-leucyl-L-leucine methyl ester-mediated killing of cytotoxic lymphocytes: dependence on a lysosomal thiol protease, dipeptidyl peptidase I, that is enriched in these cells. Proceedings of the National Academy of Sciences of the United States of America 87, 83–87 (1990).

38. Uchimoto, T. et al. Mechanism of apoptosis induced by a lysosomotropic agent, L-Leucyl-L-Leucine methyl ester. Apoptosis 4, 357–362 (1999).

39. Aits, S. et al. Sensitive detection of lysosomal membrane permeabilization by lysosomal galectin puncta assay. Autophagy 11, 1408–1424 (2015).

40. Eapen, V.V., Swarup, S., Hoyer, M.J., Paulo, J.A. & Harper, J.W. Quantitative proteomics reveals the selectivity of ubiquitin-binding autophagy receptors in the turnover of damaged lysosomes by lysophagy. Elife 10 (2021).

41. Papadopoulos, C. & Meyer, H. Detection and Clearance of Damaged Lysosomes by the Endo-Lysosomal Damage Response and Lysophagy. Curr Biol 27, R1330–r1341 (2017).

42. Kirkegaard, T. et al. Hsp70 stabilizes lysosomes and reverts Niemann-Pick disease-associated lysosomal pathology. Nature 463, 549–553 (2010).

43. Lee, J.H. et al. Faulty autolysosome acidification in Alzheimer’s disease mouse models induces autophagic build-up of Aβ in neurons, yielding senile plaques. Nature neuroscience 25, 688–701 (2022).

44. Neely Kayala, K.M., et al. Presenilin-null cells have altered two-pore calcium channel expression and lysosomal calcium: implications for lysosomal function. Brain research 1489, 8–16 (2012).

45. Bretou, M. et al. Accumulation of APP C-terminal fragments causes endolysosomal dysfunction through the dysregulation of late endosome to lysosome-ER contact sites. Developmental cell 59, 1571–1592.e1579 (2024).

46. Caramello, A. et al. Intra-cellular accumulation of amyloid is a marker of selective neuronal vulnerability in Alzheimer’s disease. medRxiv, 2023.2011.2023.23298911 (2023).

47. Nixon, R.A., Yang, D.S. & Lee, J.H. Neurodegenerative lysosomal disorders: a continuum from development to late age. Autophagy 4, 590–599 (2008).

48. Faizi, M. et al. Thy1-hAPP(Lond/Swe+) mouse model of Alzheimer’s disease displays broad behavioral deficits in sensorimotor, cognitive and social function. Brain Behav 2, 142–154 (2012).

49. Heneka, M.T. et al. NLRP3 is activated in Alzheimer’s disease and contributes to pathology in APP/PS1 mice. Nature 493, 674–678 (2013).

50. Vest, R.T. et al. Small molecule C381 targets the lysosome to reduce inflammation and ameliorate disease in models of neurodegeneration. Proceedings of the National Academy of Sciences of the United States of America 119, e2121609119 (2022).

51. Rentzos, M. et al. IL-15 is elevated in cerebrospinal fluid of patients with Alzheimer’s disease and frontotemporal dementia. Journal of geriatric psychiatry and neurology 19, 114–117 (2006).

52. Moreau, D. et al. Drug-induced increase in lysobisphosphatidic acid reduces the cholesterol overload in Niemann-Pick type C cells and mice. EMBO reports 20, e47055 (2019).

53. Benyair, R. et al. Upregulation of the ESCRT pathway and multivesicular bodies accelerates degradation of proteins associated with neurodegeneration. Autophagy Rep 2 (2023).

54. Rowe, R.G. & Daley, G.Q. Induced pluripotent stem cells in disease modelling and drug discovery. Nat Rev Genet 20, 377–388 (2019).

55. Hansson, J. et al. Highly coordinated proteome dynamics during reprogramming of somatic cells to pluripotency. Cell reports 2, 1579–1592 (2012).

56. Miller, J.D. et al. Human iPSC-based modeling of late-onset disease via progerin-induced aging. Cell Stem Cell 13, 691–705 (2013).

57. Huh, C.J. et al. Maintenance of age in human neurons generated by microRNA-based neuronal conversion of fibroblasts. Elife 5 (2016).

58. Herdy, J. et al. Chemical modulation of transcriptionally enriched signaling pathways to optimize the conversion of fibroblasts into neurons. Elife 8 (2019).

59. Wingo, A.P. et al. Integrating human brain proteomes with genome-wide association data implicates new proteins in Alzheimer’s disease pathogenesis. Nature genetics 53, 143–146 (2021).

60. Rubinsztein, D.C., Mariño, G. & Kroemer, G. Autophagy and aging. Cell 146, 682–695 (2011).

61. Cataldo, A.M. & Nixon, R.A. Enzymatically active lysosomal proteases are associated with amyloid deposits in Alzheimer brain. Proceedings of the National Academy of Sciences of the United States of America 87, 3861–3865 (1990).

62. Nixon, R.A. Autophagy, amyloidogenesis and Alzheimer disease. J Cell Sci 120, 4081–4091 (2007).

63. Gowrishankar, S. et al. Massive accumulation of luminal protease-deficient axonal lysosomes at Alzheimer’s disease amyloid plaques. Proceedings of the National Academy of Sciences of the United States of America 112, E3699–3708 (2015).

64. Sanyal, A., et al. Constitutive Endolysosomal Perforation in Neurons allows Induction of α-Synuclein Aggregation by Internalized Pre-Formed Fibrils. bioRxiv (2024).

65. Rose, K. et al. Tau fibrils induce nanoscale membrane damage and nucleate cytosolic tau at lysosomes. Proceedings of the National Academy of Sciences of the United States of America 121, e2315690121 (2024).

66. Van Acker, Z.P. et al. Phospholipase D3 degrades mitochondrial DNA to regulate nucleotide signaling and APP metabolism. Nature communications 14, 2847 (2023).

67. Coen, K. et al. Lysosomal calcium homeostasis defects, not proton pump defects, cause endo-lysosomal dysfunction in PSEN-deficient cells. The Journal of cell biology 198, 23–35 (2012).

68. Colacurcio, D.J. & Nixon, R.A. Disorders of lysosomal acidification-The emerging role of v-ATPase in aging and neurodegenerative disease. Ageing research reviews 32, 75–88 (2016).

69. Nixon, R.A. Amyloid precursor protein and endosomal-lysosomal dysfunction in Alzheimer’s disease: inseparable partners in a multifactorial disease. Faseb j 31, 2729–2743 (2017).

70. Willén, K. et al. Aβ accumulation causes MVB enlargement and is modelled by dominant negative VPS4A. Molecular neurodegeneration 12, 61 (2017).

71. Kwart, D. et al. A Large Panel of Isogenic APP and PSEN1 Mutant Human iPSC Neurons Reveals Shared Endosomal Abnormalities Mediated by APP β-CTFs, Not Aβ. Neuron 104, 256–270.e255 (2019).

72. Kaushik, S. et al. Autophagy and the hallmarks of aging. Ageing research reviews 72, 101468 (2021).

73. Stavoe, A.K.H. & Holzbaur, E.L.F. Neuronal autophagy declines substantially with age and is rescued by overexpression of WIPI2. Autophagy 16, 371–372 (2020).

74. Niekamp, P. et al. Ca(2+)-activated sphingomyelin scrambling and turnover mediate ESCRT-independent lysosomal repair. Nature communications 13, 1875 (2022).

75. Zhu, S.Y. et al. Lysosomal quality control of cell fate: a novel therapeutic target for human diseases. Cell death & disease 11, 817 (2020).

76. Bonet-Ponce, L. et al. LRRK2 mediates tubulation and vesicle sorting from lysosomes. Sci Adv 6 (2020).

77. Ebstrup, M.L. et al. Annexin A7 mediates lysosome repair independently of ESCRT-III. Front Cell Dev Biol 11, 1211498 (2023).

78. Funk, K.E., Mrak, R.E. & Kuret, J. Granulovacuolar degeneration (GVD) bodies of Alzheimer’s disease (AD) resemble late-stage autophagic organelles. Neuropathology and applied neurobiology 37, 295–306 (2011).

79. Hondius, D.C. et al. The proteome of granulovacuolar degeneration and neurofibrillary tangles in Alzheimer’s disease. Acta neuropathologica 141, 341–358 (2021).

80. Gavin, A.L. et al. PLD3 and PLD4 are single-stranded acid exonucleases that regulate endosomal nucleic-acid sensing. Nat Immunol 19, 942–953 (2018).

81. Satoh, J. et al. PLD3 is accumulated on neuritic plaques in Alzheimer’s disease brains. Alzheimer’s research & therapy 6, 70 (2014).

82. Deretic, V. Autophagy in inflammation, infection, and immunometabolism. Immunity 54, 437–453 (2021).

83. Zhang, R., Kang, R. & Tang, D. The STING1 network regulates autophagy and cell death. Signal Transduct Target Ther 6, 208 (2021).

84. Wang, D. et al. The role of NLRP3-CASP1 in inflammasome-mediated neuroinflammation and autophagy dysfunction in manganese-induced, hippocampal-dependent impairment of learning and memory ability. Autophagy 13, 914–927 (2017).

85. Hipp, M.S., Kasturi, P. & Hartl, F.U. The proteostasis network and its decline in ageing. Nature reviews. Molecular cell biology 20, 421–435 (2019).

86. Stein, K.C., Morales-Polanco, F., van der Lienden, J., Rainbolt, T.K. & Frydman, J. Ageing exacerbates ribosome pausing to disrupt cotranslational proteostasis. Nature 601, 637–642 (2022).

## References (Methods)

87. Tanabe, K. et al. Transdifferentiation of human adult peripheral blood T cells into neurons. Proceedings of the National Academy of Sciences of the United States of America (2018).

88. Rockenstein, E., Mallory, M., Mante, M., Sisk, A. & Masliaha, E. Early formation of mature amyloid-beta protein deposits in a mutant APP transgenic model depends on levels of Abeta(1-42). Journal of neuroscience research 66, 573–582 (2001).

89. Nguyen, A.T. et al. UBE2O remodels the proteome during terminal erythroid differentiation. Science (New York, N.Y.) 357 (2017).

90. Cao, C. et al. Maternal Iron Deficiency Modulates Placental Transcriptome and Proteome in Mid-Gestation of Mouse Pregnancy. The Journal of nutrition 151, 1073–1083 (2021).

91. Liu, X., Li, J., Gygi, S.P. & Paulo, J.A. Profiling Yeast Deletion Strains Using Sample Multiplexing and Network-Based Analyses. Journal of proteome research 21, 1525–1536 (2022).

92. Rappsilber, J., Ishihama, Y. & Mann, M. Stop and go extraction tips for matrix-assisted laser desorption/ionization, nanoelectrospray, and LC/MS sample pretreatment in proteomics. Analytical chemistry 75, 663–670 (2003).

93. Schweppe, D.K. et al. Characterization and Optimization of Multiplexed Quantitative Analyses Using High-Field Asymmetric-Waveform Ion Mobility Mass Spectrometry. Analytical chemistry 91, 4010–4016 (2019).

94. Schweppe, D.K., Rusin, S.F., Gygi, S.P. & Paulo, J.A. Optimized Workflow for Multiplexed Phosphorylation Analysis of TMT-Labeled Peptides Using High-Field Asymmetric Waveform Ion Mobility Spectrometry. Journal of proteome research 19, 554–560 (2020).

95. Schweppe, D.K. et al. Full-Featured, Real-Time Database Searching Platform Enables Fast and Accurate Multiplexed Quantitative Proteomics. Journal of proteome research 19, 2026–2034 (2020).

96. Eng, J.K., McCormack, A.L. & Yates, J.R. An approach to correlate tandem mass spectral data of peptides with amino acid sequences in a protein database. J Am Soc Mass Spectrom 5, 976–989 (1994).

97. Eng, J.K., Fischer, B., Grossmann, J. & Maccoss, M.J. A fast SEQUEST cross correlation algorithm. Journal of proteome research 7, 4598–4602 (2008).

98. Eng, J.K., Jahan, T.A. & Hoopmann, M.R. Comet: an open-source MS/MS sequence database search tool. Proteomics 13, 22–24 (2013).

99. Huttlin, E.L. et al. A tissue-specific atlas of mouse protein phosphorylation and expression. Cell 143, 1174–1189 (2010).

100. Vizcaíno, J.A. et al. ProteomeXchange provides globally coordinated proteomics data submission and dissemination. Nature biotechnology 32, 223–226 (2014).

101. Villeda, S.A. et al. Young blood reverses age-related impairments in cognitive function and synaptic plasticity in mice. Nature medicine 20, 659–663 (2014).

102. Gold, L. et al. Aptamer-based multiplexed proteomic technology for biomarker discovery. PloS one 5, e15004 (2010).

103. Kim, C.H. et al. Stability and reproducibility of proteomic profiles measured with an aptamer-based platform. Scientific reports 8, 8382 (2018).

