## Extended Data Figure Legends for "Proteostasis and lysosomal repair deficits in transdifferentiated neurons of Alzheimer’s disease"

| | | |
|---|---|---|
| Extended Data Fig. 1 | Proteostasis signatures of aging and Alzheimer's disease (AD) are present in human adult fibroblasts. | **(a)** Representative images of endogenous ubiquitin and p62/SQSTM1 in young fibroblasts treated with DMSO control (Ctrl), proteasome inhibitor Bortezomib (BTZ, 25 nM) or lysosome inhibitor Chloroquine (CQ, 25 μM) for 24 hr (scale bar: 40 μm). Insert: higher magnification view of ubiquitin and p62/SQSTM1 (scale bar: 20 μm). **(b)** Immunofluorescence (IF) quantification of proteostasis markers shown in panel (a) for changes in ubiquitin and p62/SQSTM1 by the treatment of BTZ and CQ, respectively, in young (Y), aged (A) and aged/sAD (sAD) fibroblasts. Data represent as fold changes in ubiquitin and p62/SQSTM1 levels (% of area per image field) relative to young fibroblasts treated with DMSO Ctrl. Ubiquitin: Ctrl: *n* = 230 (young), 215 (aged) and 162 (aged/sAD) cells; BTZ: *n* = 124 (young), 142 (aged) and 131 (aged/sAD) cells; CQ: *n* = 152 (young), 131 (aged) and 134 (aged/sAD) cells. p62/SQSTM1: Ctrl: *n* = 140 (young), 117 (aged) and 138 (aged/sAD) cells; BTZ: *n* = 155 (young), 130 (aged) and 143 (aged/sAD) cells; CQ: *n* = 133 (young), 150 (aged) and 137 (aged/sAD) cells.<br>In panel **b**, the boxes show median and $1^{st}$ and $3^{rd}$ quartile and the whiskers extending 1.5 times the interquartile range from the boxes. Data show box-and-whisker plots of three to five independent experiments and cells from three independent healthy control (HC) and AD donors. Statistical analysis is performed using Two-Way ANOVA followed by Bonferroni post-hoc analysis. **P < 0.01 and ***P < 0.001. Source numerical data are available in Source data. |
| Extended Data Fig. 2 | Cortical neurons are generated directly from human adult fibroblasts using a combinatorial | **(a)** Transdifferentiation scheme using transcription factors *Brn2*, *Ascl1*, *Myt1l*, and *Ngn2* (herein BAMN factors) that induces a global transcriptional change in fibroblasts to |

| | | |
|---|---|---|
| | transcription-factor and small-molecule protocol. | constitute a new neuronal cell state. Fibroblasts are plated one day before transdifferentiation. On Day 0, cells are infected with lentiviruses expressing BAMN factors and harvested four days after doxycycline-induction. Transduced cells expressing PSA-NCAM are magnetically isolated and replated to vitronectin/laminin-coated plates on Day 4 and then switched to reprogramming medium (DMEM/F12/Neurobasal) supplemented with small molecules 5 μM Forskolin, 10 μM SB 431542, 2 μM Dorsomorphin and 2 μM XAV939 on Day 5. 10 ng/mL BDNF and NT-3 are added into the reprogramming medium after one week. Cells are switched to maturation medium (Neuronal) on Day 20 and used for experiments after Day 35. **(b)** Fibroblasts are infected with lentiviruses expressing BAMN factors and GFP and monitored for morphological changes after lentiviral induction. Transduced cells undergo sorting based on PSA-NCAM+ gate in flow cytometry on Day 4. Scale bar: 200 μm. **(c,d)** IF images of neuronal markers in tNeurons at PID 35. Quantification of the percentage of cells positive for both NeuN and Tuj1 (c) and the numbers of Syn-1 puncta in the soma (d). Tuj1/DAPI, MAP2/DAPI & MAP2/Tuj1: *n* = 1208 (young), 678 (aged) and 789 (aged/sAD) cells; NeuN/DAPI & NeuN/Tuj1: *n* = 822 (young), 579 (aged) and 759 (aged/sAD) cells. Syn-1: *n* = 40 (young), 42 (aged) and 41 (aged/sAD) cells. Scale bar: 100 μm (c) and 20 μm (d). In panel **c** and **d**, the boxes show median and 1$^{st}$ and 3$^{rd}$ quartile and the whiskers extending 1.5 times the interquartile range from the boxes. Data show box-and-whisker plots of two to four independent experiments and cells from independent HC and AD patients. Statistical analysis is performed using One-Way ANOVA followed by Bonferroni post-hoc analysis. *P < 0.05 and ***P < 0.001. |

| | | |
|---|---|---|
| | | Source numerical data are available in Source data. |
| Extended Data Fig. 3 | Human tNeurons derived from sAD patients exhibit disease-related protein pathologies. | **(a)** IF images and quantification of proteostasis- and disease-associated protein markers, including p62/SQSTM1, ubiquitin, total Aβ and the toxic isoform Aβ42, hyper-phosphorylated tau (pTau) and TDP-43 (pTDP-43) and nuclear-to-cytoplasmic (N-to-C) ratio of TDP-43 in tNeurons shown as full images for Fig.1g. Cells are co-stained with Tuj1. White dash line represents the nuclear region (N). p62/SQSTM1: $n$ = 62 (young), 62 (aged) and 66 (aged/sAD) cells; ubiquitin: $n$ = 46 (young), 53 (aged) and 66 (aged/sAD) cells; total Aβ: $n$ = 111 (young), 106 (aged) and 114 (aged/sAD) cells; Aβ42: $n$ = 103 (young), 95 (aged) and 93 (aged/sAD) cells; pTau: $n$ = 91 (young), 89 (aged) and 114 (aged/sAD) cells; pTDP-43: $n$ = 91 (young), 74 (aged) and 95 (aged/sAD) cells; N-to-C ratio of TDP-43: $n$ = 85 (young), 86 (aged) and 103 (aged/sAD) cells. Scale bar: 20 μm. **(b)** IF staining and quantification of small heat shock protein HspB1 in tNeurons. $n$ = 80 (young), 111 (aged) and 134 (aged/sAD) cells. Scale bar: 20 μm. **(c)** Sandwich ELISA assay for detecting endogenous Aβ42 in total cell lysates of tNeurons. The values are revealed by a fold change relative to young tNeurons. $n$ = 4 (young), 4 (aged) and 4 (aged/sAD) independent replicates. **(d)** IF staining of pTau and pTDP-43 with p62/SQSTM1 and ubiquitin in aged/sAD tNeurons (scale bar: 20 μm). Cells are co-stained with Tuj1 and DAPI. White dash line represents the nuclear region (N). Insert: higher magnification view of protein colocalization within individual neuron (scale bar: 5 μm). Arrowhead: protein colocalization. In panel **a**, **b** and **c**, the boxes show median and 1$^{st}$ and 3$^{rd}$ quartile and the whiskers extending 1.5 times the interquartile range from the boxes. Data show box-and-whisker plots of two to three |

| | | |
|---|---|---|
| | | independent experiments and cells from independent HC and AD patients. Statistical analysis is performed using One-Way ANOVA followed by Bonferroni post-hoc analysis. *P < 0.05, **P < 0.01 and ***P < 0.001. Source numerical data are available in Source data. |
| Extended Data Fig. 4 | Quantitative proteomic analysis of human tNeurons identifies proteins and pathways associated with aging and sAD. | **(a)** Heatmap of the differentially expressed proteins across tNeurons from young, aged, aged/sAD and fAD-PSEN1 donors. **(b)** Top-ranked proteins (rows) changing with aging and sAD (columns) based on log2-fold change (log2-FC). **(c)** Gene ontology (GO) analysis of the differentially expressed proteins between young, aged and aged/sAD tNeurons. Circle sizes reflect the number (#) of proteins. **(d)** Network analysis of tNeuron proteomes, including proteins in the top-ranked pathways associated with young and sAD, revealed by GeneMANIA and GO. Comparison between healthy young (*n* = 3 individuals) and aged/sAD tNeurons (*n* = 6 individuals) at PID 40. Colored circles represent the enrichment of identified proteins revealing by log2-FC: increase in red and decrease in blue. **(e)** Venn diagram reveals an overlap of differentially expressed proteins between tNeurons from aged and young donors, and between tNeurons from aged/sAD and young donors. The shared hits and assigned GO terms are showed in light green color. Interaction network for the shared hits that are either increased or decreased in aged and aged/sAD tNeurons as compared with young tNeurons. Each node representing a single protein that is divided to reflect the individual change for aged vs. young (left) and aged/sAD vs. young (right). Protein abundance increases are shown in red, and decreases shown in blue. |
| Extended Data Fig. 5 | Quantitative proteomic and molecular analysis of tNeurons identify signatures of | **(a)** IF images and quantification of proteostasis- and disease-associated protein markers in tNeurons from healthy young donors as well as AD patients with aged/sAD and fAD-PSEN1. |

| | | |
|---|---|---|
| | pathogenic mutations in the *PSEN1* gene in AD patient neurons. | White dash line represents the nuclear region (N). p62/SQSTM1: *n* = 62 (young), 66 (aged/sAD) and 49 (fAD-PSEN1) cells; ubiquitin: *n* = 46 (young), 66 (aged/sAD) and 50 (fAD-PSEN1) cells; total Aβ: *n* = 111 (young), 114 (aged/sAD) and 57 (fAD-PSEN1) cells; Aβ42: *n* = 103 (young), 93 (aged/sAD) and 68 (fAD-PSEN1) cells; pTau: *n* = 91 (young), 114 (aged/sAD) and 107 (fAD-PSEN1) cells; pTDP-43: *n* = 91 (young), 95 (aged/sAD) and 82 (fAD-PSEN1) cells; N-to-C ratio of TDP-43: *n* = 85 (young), 103 (aged/sAD) and 109 (fAD-PSEN1) cells. Scale bar: 20 µm. **(b)** Detection of endogenous Aβ42 in total cell lysates of tNeurons using sandwich ELISA assay. The values are revealed by a fold change relative to young tNeurons. *n* = 4 (young), 4 (aged/sAD) and 4 (fAD-PSEN1) independent replicates. **(c)** IF quantification of small heat shock protein HspB1 in young, aged/sAD and fAD-PSEN1 tNeurons. *n* = 80 (young), 134 (aged/sAD) and 136 (fAD-PSEN1) cells. **(d)** Top-ranked proteins (rows) changing with aged/sAD and fAD-PSEN1 (columns) based on log2-FC. **(e)** Network analysis of tNeuron proteins in the top-ranked pathways associated with aged/sAD and fAD-PSEN1. Comparison between aged/sAD (*n* = 6 individuals) and fAD-PSEN1 (*n* = 2 individuals) at PID 40. Colored circles represent the enrichment of identified proteins revealing by log2-FC: increase in red and decrease in blue. **(f)** GO analysis of the differentially expressed proteins across aged/sAD and fAD-PSEN1 tNeurons. Circle sizes reflect the number (#) of proteins. **(g)** Lists of top-ranked proteins (rows) changing with fAD-PSEN1 and young (columns) based on log2-FC. **(h)** Network analysis and differential expression of proteins detected in tNeurons from healthy young donors (*n* = 3 individuals) and patients with fAD-PSEN1 (*n* = 2 individuals) at PID 40. The top-ranked pathways for fAD-PSEN1 proteome are analyzed using GO databases. Colored circles |

| | | |
|---|---|---|
| | | represent the enrichment of identified proteins revealing by log2-FC: increase in red and decrease in blue. **(i)** GO analysis of the differentially expressed proteins across fAD-PSEN1 and young tNeurons. Circle sizes reflect the number (#) of proteins. In panel **a**, **b** and **c**, the boxes show median and $1^{st}$ and $3^{rd}$ quartile and the whiskers extending 1.5 times the interquartile range from the boxes. Data show box-and-whisker plots of two to three independent experiments and cells from independent HC and AD patients. Statistical analysis is performed using One-Way ANOVA followed by Bonferroni post-hoc analysis. *P < 0.05, **P < 0.01 and ***P < 0.001. Source numerical data are available in Source data. |
| Extended Data Fig. 6 | Aberrant response to lysosomal damage at basal conditions and during LLOME treatment are prominent in aged/sAD tNeurons. | **(a)** Schematic for describing the ESCRT- and Galectin-mediated lysosomal quality control (LQC) machinery. When undergoing lysosomal damage stress, lysosomal membrane is subjected to rupture. To avoid the leakage of lysosomal contents and activation of cell death pathways, the compromised lysosomes recruit ESCRT proteins (ESCRTs) to repair the small membrane wounds and Galectins to target the massively damaged lysosomes for lysophagy. **(b)** Representative images for immunostaining of CHMP2B (magenta), LAMP2 (green) and Tuj1 (blue) in young, aged and aged/sAD tNeurons, related to Fig. 3c. Arrowhead: high-density accumulations of CHMP2B in the vicinity of plasma membrane (solid) and within neurites (hollow). Scale bar: 10 μm. **(c)** Representative images for immunostaining of Galectin-3 (magenta), LAMP2 (green) and Tuj1 (blue) in young, aged and aged/sAD tNeurons, related to Fig. 3c. Scale bar: 10 μm. **(d)** Measurement of spontaneous cell death, indicated by apoptosis markers (e.g. active Caspase-3/7), in young, aged, aged/sAD and fAD-PSEN1 tNeurons during |

| | | |
|---|---|---|
| | | *in vitro* culture at PID 35, 38 and 42. PID 35: *n* = 162 (young), 158 (aged), 148 (aged/sAD) and 136 (fAD-PSEN1) cells; PID 38: *n* = 163 (young), 159 (aged), 147 (aged/sAD) and 156 (fAD-PSEN1) cells; PID 42: *n* = 159 (young), 151 (aged), 142 (aged/sAD) and 145 (fAD-PSEN1) cells. **(e)** Immunostaining of TDP-43 and Hsp70 (magenta), LAMP1 (green) and Tuj1 (blue) during DMSO Ctrl or 0.25 mM LLOME treatment for 30 min in young, aged, aged/sAD tNeurons at PID 35 (scale bar: 10 µm). Insert: higher magnification view of protein colocalization within individual neuron (scale bar: 2 µm). Arrowhead: protein colocalization. **(f)** Changes in lysosomal acidification by detecting the intensity of preloaded pH-sensitive FITC-conjugated Dextran in tNeurons at basal conditions and during 0.25 mM LLOME treatment for 30 min. Ctrl: *n* = 145 (young), 135 (aged) and 148 (aged/sAD) cells; LLOME: *n* = 142 (young), 137 (aged) and 155 (aged/sAD) cells. **(g)** IF analysis of colocalization of APP-CTF with LAMP1 in aged/sAD tNeurons (scale bar: 10 µm). Insert: higher magnification view of APP-CTF and LAMP1 (scale bar: 2 µm). **(h)** IF analysis of colocalization of Aβ42 with LC3B in aged/sAD tNeurons (scale bar: 10 µm). Insert: higher magnification view of Aβ42 and LC3B (scale bar: 2 µm). In panel **d** and **f**, the boxes show median and 1st and 3rd quartile and the whiskers extending 1.5 times the interquartile range from the boxes. Data show box-and-whisker plots of three independent experiments and cell lines from independent HC and AD donors. Statistical analysis is performed using Two-Way ANOVA followed by Bonferroni post-hoc analysis. Source numerical data are available in Source data. |
| Extended Data Fig. 7 | Abnormal lysosomal damage response links to defective organelle and calcium | **(a)** Quantification of numbers of ESCRT-III CHMP2B and Galectin-3 puncta in the cell body of young, aged/sAD and fAD-PSEN1 tNeurons in the absence of lysosomal damage insults. |

| | | |
|---|---|---|
| | homeostasis in aged/sAD and fAD-PSEN1 tNeurons. | tNeurons: CHMP2B: *n* = 117 (young), 97 (aged/sAD) and 104 (fAD-PSEN1) cells; Galectin-3: *n* = 111 (young), 95 (aged/sAD) and 104 (fAD-PSEN1) cells. **(b)** Quantification of numbers of ESCRT-0 HGS puncta in the cell body of young, aged/sAD and fAD-PSEN1 tNeurons during lysosomal damage insults mediated by LLOME treatment, and recovery from lysosomal damage after LLOME washout for up to 8 hr. The half-life ($t_{1/2}$) represents the time required for lysosomal repair. Young: *n* = 108 (Ctrl), 107 (LLOME), 111 (Washout 1 hr), 118 (Washout 2 hr) and 90 (Washout 8 hr) cells; Aged/sAD: *n* = 115 (Ctrl), 115 (LLOME), 117 (Washout 1 hr), 109 (Washout 2 hr) and 79 (Washout 8 hr) cells; fAD-PSEN1: *n* = 117 (Ctrl), 117 (LLOME), 113 (Washout 1 hr), 116 (Washout 2 hr) and 85 (Washout 8 hr) cells. **(c)** Changes in lysosomal acidification revealed by the intensity of preloaded pH-sensitive FITC-conjugated Dextran in young, aged/sAD and fAD-PSEN1 tNeurons at basal conditions or during 0.25 mM LLOME treatment for 30 min. Ctrl: *n* = 145 (young), 148 (aged/sAD) and 137 (fAD-PSEN1) cells; LLOME: *n* = 142 (young), 155 (aged/sAD) and 147 (fAD-PSEN1) cells. **(d)** Quantification of changes in mitochondrial membrane potential in young, aged/sAD and fAD-PSEN1 tNeurons at basal conditions and after the treatment of DMSO Ctrl, 20 μM FCCP or 0.25 mM LLOME for 30 min by analyzing TMRE intensity. The values are revealed by a fold change relative to young tNeurons treated with DMSO. *n* = 6 (young), 6 (aged/sAD) and 6 (fAD-PSEN1) independent replicates. **(e)** Live-cell imaging and analysis of lysosomal calcium revealed by Cal-520, a fluorogenic calcium-sensitive indicator, conjugated with Dextran molecules, in young, aged, aged/sAD and fAD-PSEN1 tNeurons at basal conditions. Lysosomes are labeled with Lysotracker Red DND-99 (scale bar: 40 μm). *n* = 163 (young), 156 (aged), 187 |

| | | |
|---|---|---|
| | | (aged/sAD) and 139 (fAD-PSEN1) cells. Insert: higher magnification view of Cal-520 and Lysotracker Red DND-99 (scale bar: 5 μm). **(f)** Analysis of correlation between intra-cellular Aβ42 levels and lysosomal acidification or calcium in young, aged, aged/sAD and fAD-PSEN1 tNeurons. *n* = 9 independent replicates from three donors (lysosomal acidification) and *n* = 6 independent replicates from two donors (lysosomal calcium) from three independent experiments. Black solid line represents the fitted linear correlation. Coefficient of Discrimination ($R^2$) is calculated using Pearson's correlation. In panel **a**, **b**, **c**, and **e**, the boxes show median and 1$^{st}$ and 3$^{rd}$ quartile and the whiskers extending 1.5 times the interquartile range from the boxes. In panel **d**, data are displayed as mean ± SD. Data show mean and SD or box-and-whisker plots of two to three independent experiments and three cell lines from independent HC and AD donors. Statistical analysis is performed using One-Way ANOVA (**a**, and **e**) or Two-Way ANOVA (**b**, **c** and **d**) followed by Bonferroni post-hoc analysis. *P < 0.05, **P < 0.01 and ***P < 0.001. ###P < 0.001. Source numerical data are available in Source data. |
| Extended Data Fig. 8 | Lysosomal damage correlates with aging and intrinsic disease properties in AD patient neurons. | **(a)** Analysis of correlation between intra-cellular Aβ42 levels and CHMP2B puncta numbers in young, aged, aged/sAD and fAD-PSEN1 tNeurons. **(b)** Analysis of correlation between intra-cellular total Aβ levels and Galectin-3 or CHMP2B puncta numbers in young, aged, aged/sAD and fAD-PSEN1 tNeurons. **(c)** Analysis of correlation between lysosomal acidification and Galectin-3 or CHMP2B puncta numbers in young, aged, aged/sAD and fAD-PSEN1 tNeurons. **(d)** Analysis of correlation between lysosomal calcium and Galectin-3 or CHMP2B puncta numbers in young, aged, aged/sAD and fAD-PSEN1 tNeurons. Panel **a**, **b** and **c**: *n* = 9 |

| | | |
|---|---|---|
| | | independent replicates from three donors and three experiments; Panel **d**: *n* = 6 independent replicates from two donors and three experiments. Black solid line represents the fitted linear correlation. Coefficient of Discrimination ($R^2$) is calculated using Pearson's correlation. Source numerical data are available in Source data. |
| Extended Data Fig. 9 | Lysosomal damage markers are elevated in the cortex of old mice and associated with neuropathological signatures of AD in the cortex of APP transgenic mice. | **(a)** Experimental schematic for wild-type young (3-month old) and old (20 to 24-month old) mice. **(b)** Representative images and IF quantification of Galectin-3 immunoreactivity and lysosomal size based on LAMP1 signals in neurons of wild-type young and old mice. Galectin-3: *n* = 3 mice; Lysosomal size: *n* = 4 mice; Scale bar: 50 µm. **(c)** Experimental schematic for non-transgenic wild-type mice (NTg) and transgenic mice expressing mutant human APP with the Swedish (K670N/M671L) and London (V717I) mutations ($APP^{Lon/Swe}$) (3 to 6-month old). **(d)** IF staining and quantification of Aβ42, LAMP1, CHMP2B and Galectin-3 in the neocortex of NTg and $APP^{Lon/Swe}$ transgenic mice, related to Fig. 4b. Extra-cellular and intra-neuronal colocalization of CHPM2B with LAMP1 in the brain of $APP^{Lon/Swe}$ mice. The brain tissue is co-stained with MAP2 and Hoechst (scale bar: 50 µm). Aβ42: *n* = 8 (NTg) and 13 ($APP^{Lon/Swe}$) mice; LAMP1: *n* = 8 (NTg) and 13 ($APP^{Lon/Swe}$) mice; CHMP2B: *n* = 10 (NTg) and 11 ($APP^{Lon/Swe}$) mice; Galectin-3: *n* = 10 (NTg) and 12 ($APP^{Lon/Swe}$) mice. Insert: higher magnification view of colocalization between CHMP2B, Galectin-3, LAMP1 and Aβ42 (scale bar: 10 µm). Arrowhead: CHMP2B and LAMP1 colocalization in an individual neuron. **(e)** Immunostaining of Galectin-3 (magenta), LAMP1 (cyan) and MAP2 (grey) counterstained with Hoechst (yellow) in the cortex of NTg and $APP^{Lon/Swe}$ mice (scale bar: 50 µm). Inset: higher magnification view of LAMP1-positive inclusions (scale bar: 50 µm). **(f)** |

| | | |
|---|---|---|
| | | Immunostaining of Hsp70 (magenta), LAMP1 (cyan) and MAP2 (grey) counterstained with Hoechst (yellow) in the cortex of NTg and $APP^{Lon/Swe}$ mice (scale bar: 50 μm). Inset: higher magnification view of LAMP1-positive inclusions (scale bar: 50 μm). Panel **b** and **d** are displayed as mean ± SD from three to four independent experiments in aging and AD mice model. Statistical analysis is performed using two-sided Student's *t*-test. *P < 0.05, **P < 0.01 and ***P < 0.001. Source numerical data are available in Source data. |
| Extended Data Fig. 10 | AD patients show prominent lysosomal damage phenotypes in brain tissue and elevated inflammatory factor secretion in CSF samples. | **(a-c)** Immunostaining of LAMP2 (cyan) and MAP2 (grey) along with CHMP2B (a), Galectin-3 (b) or Aβ(6E10) (c) (magenta), in the post-mortem cerebral cortex of HC and AD donors, related to Fig. 4c (scale bar: 20 μm). #: indicates images acquired from different individuals. Inset: higher magnification view of LAMP2-positive inclusions within individual neuron (scale bar: 10 μm). **(d)** IF quantification of LAMP2, CHMP2B, Galectin-3 and Aβ(6E10) in the post-mortem cerebral cortex of HC and AD donors. HC: *n* = 4; AD: *n* = 4 individuals. **(e)** Secreted inflammatory factors by human tNeurons are detected in human brain cells. Single-cell transcriptomic analysis of selected cytokines and chemokines from Fig. 5d,e in major cell types of human brain, including neuron, astrocyte, microglia and oligodendrocyte. The transcript expression scores (H: high; M: medium; L: low) is determined by the expression level of each transcript in public datasets retrieved from the Human Cell Atlas and Allen Brain Map. **(f)** Quantification of changes in the levels of inflammatory panel biomarkers measured in human CSF from HC and AD donors using the SOMAScan assay platform. HC: *n* = 50; AD: *n* = 29 individuals. In panel **d**, IF data are displayed as mean ± SD. In panel **f**, the boxes show median |

| | | |
|---|---|---|
| | | and 1st and 3rd quartile and the whiskers extending 1.5 times the interquartile range from the boxes. Statistical analysis is performed using two-sided Student's *t*-test. *P < 0.05, **P < 0.01. Source numerical data are available in Source data. |
