## Supplementary Figures for "Proteostasis and lysosomal repair deficits in transdifferentiated neurons of Alzheimer’s disease"

**Basal condition and chronic lysosomal damage in human tNeurons**

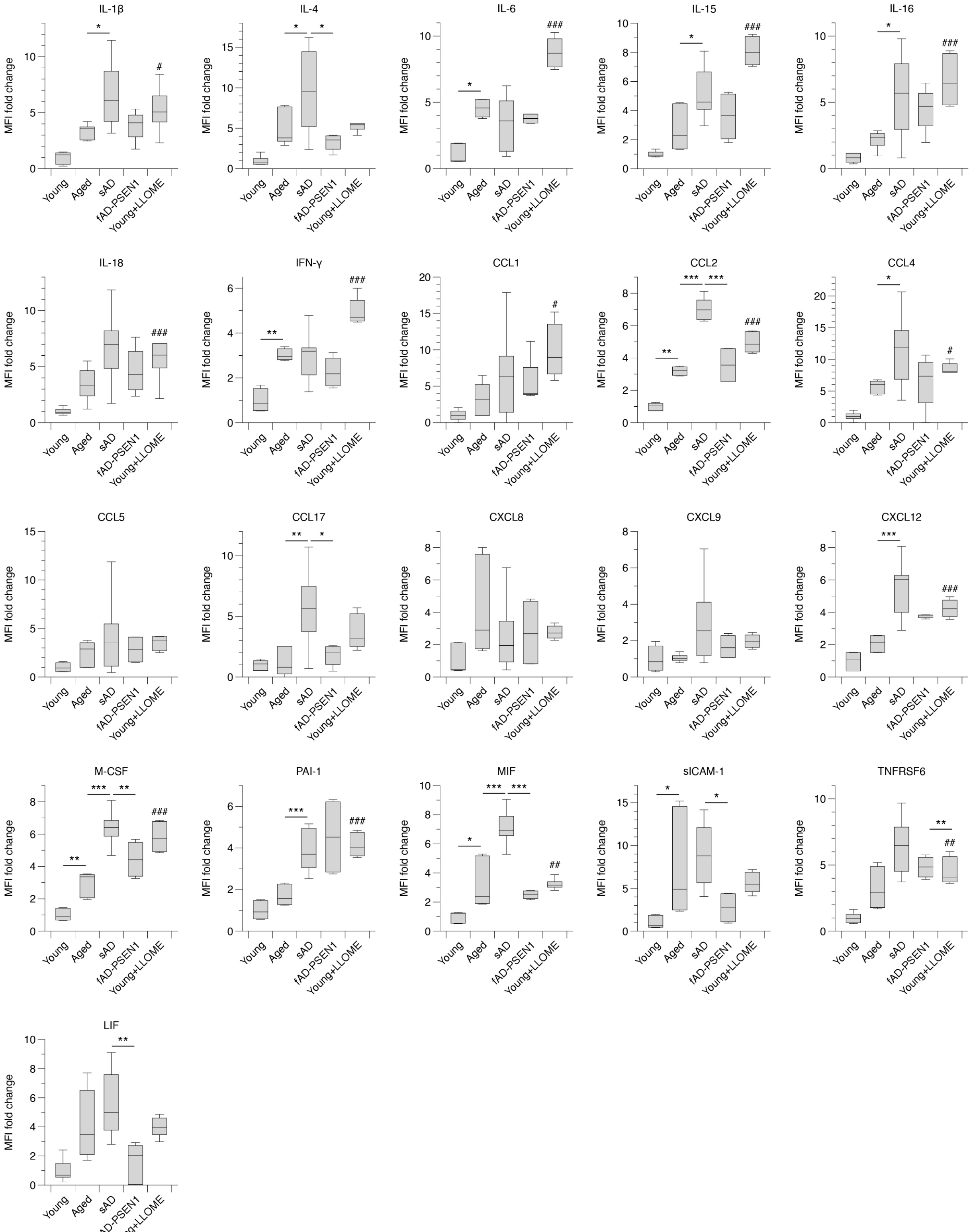

**Supplementary Fig.1: Secretion of inflammatory factors from human tNeurons is linked to AD phenotypes.**

Plots with statistical analysis of changes in inflammatory factor levels in the conditioned medium from young, aged, aged/sAD and fAD-PSEN1 tNeurons at basal conditions and young tNeurons with the treatment with 0.1 mM LLOME for 7 days, related to Fig. 5d,e.

Data show box-and-whisker plots of inflammatory profiling conducted in technical replicates and 2 to 6 cell lines in tNeurons. Box-and-whisker plots depict 5th, 25th, 50, 75th and 95th percentiles. Differences between groups are compared with One-Way ANOVA followed by Bonferroni post-hoc analysis. *$P < 0.05$, **$P < 0.01$ and ***$P < 0.001$. #$P < 0.05$, ##$P < 0.01$ and ###$P < 0.001$ as compared between young tNeurons with and without LLOME treatment.

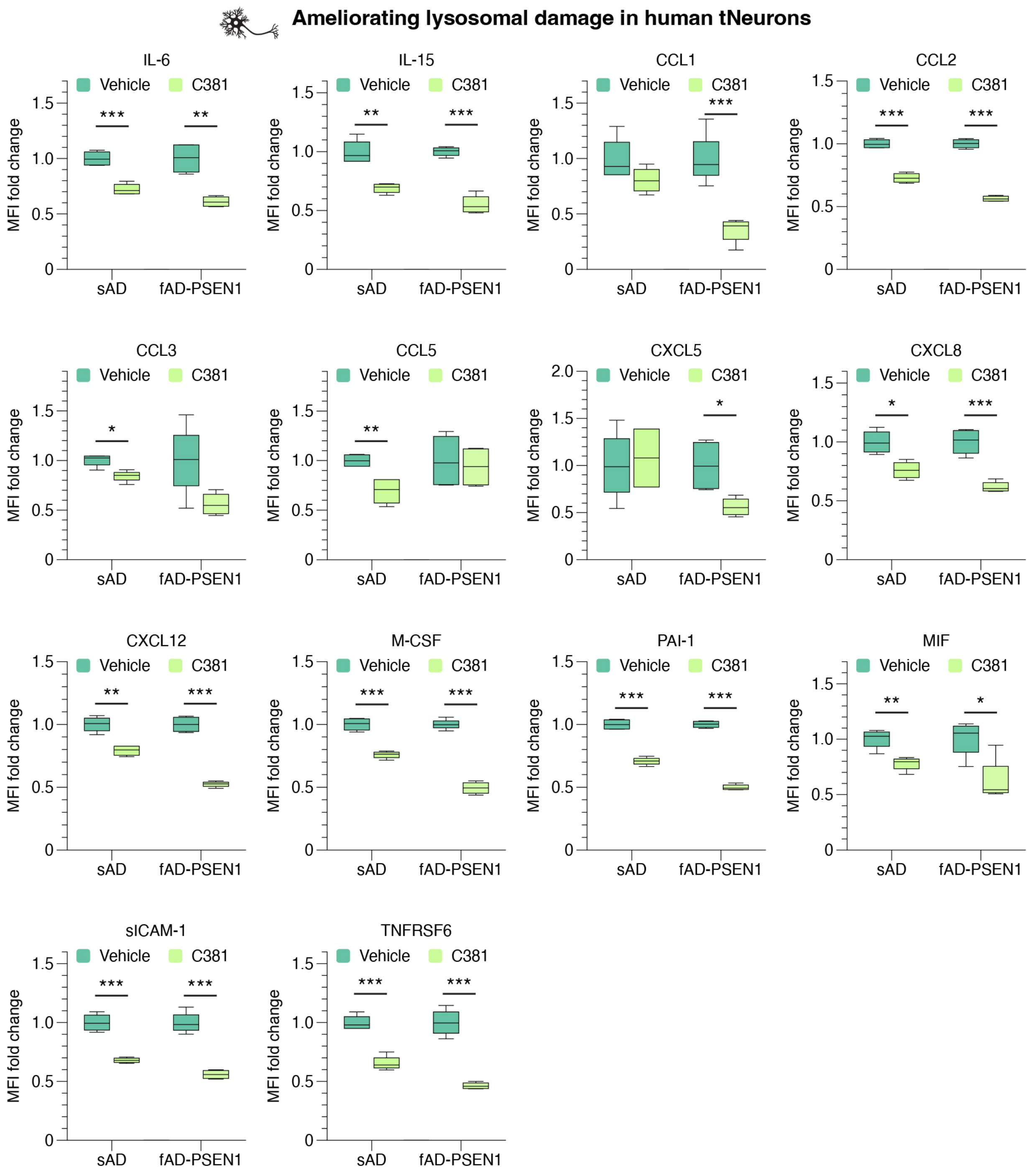


**Supplementary Fig.2: Secretion of inflammatory factors from human tNeurons can be reduced by ameliorating lysosomal deficits.**

Plots with statistical analysis of changes in inflammatory factor levels in the conditioned medium from aged/sAD and fAD-PSEN1 tNeurons with the treatment of DMSO (vehicle) or 3.1 μM C381 for 7 days as shown in Fig. 5g.

Data show box-and-whisker plots of inflammatory profiling conducted in technical replicates and 2 cell lines in tNeurons. Box-and-whisker plots depict 5th, 25th, 50, 75th and 95th percentiles. Differences between groups are compared with two-sided Student's *t*-test. *P < 0.05, **P < 0.01 and ***P < 0.001.
